# Coupling between acid-sensing ion channel 1a and the monocarboxylate transporter family shapes cellular pH response

**DOI:** 10.1101/2025.06.30.651016

**Authors:** M.H. Poulsen, N. Ritter, S. Maurya, F. Xue, S. Haider, S.G. Usher, S.A. Heusser, H.C. Chua, J. Fisker-Andersen, S. Kickinger, D. Schramek, P. Wellendorph, A. Lundby, S.A. Pless

## Abstract

Proton-mediated activation of acid-sensing ion channel 1a (ASIC1a) leads to influx of sodium and, to a lesser extent, calcium, followed by rapid and complete desensitization. However, various ASIC1a-expressing cell types display atypical proton-induced currents characterized by altered proton sensitivity and slow and incomplete desensitization. The origin of this functional diversity and its physiological relevance remain unclear. Here, we show that the monocarboxylate transporter 1 (MCT1) interacts with ASIC1a, leading to subtype-specific modulation of proton sensitivity and desensitization. We delineate the molecular mechanism of the ASIC1a-MCT1 coupling and find that the presence of MCT1 is required for the distinct ASIC1a-mediated currents observed in glioblastoma cells, the most lethal brain cancer in adults. Strikingly, presence of ASIC1a together with MCT1 significantly prolongs the lifespan of mice intracranially injected with glioblastoma cells. We thus uncover the basis for altered ASIC1a function in glioblastoma and highlight the importance of deciphering the cell-type specific ASIC1a interactome.

## Introduction

Proton concentrations are tightly regulated and play crucial roles in both the central- and peripheral-nervous system by maintaining pH homeostasis, modulating synaptic transmission, and contributing to pain perception.^1^ The discovery of protons as *de facto* neuromodulators in1980 was serendipitous ^2–5^, and remained controversial until 1997, when the first receptor responsible for these proton-dependent currents were cloned and identified as an acid-sensing ion channel (ASIC).^6^ ASICs are trimeric cation channels that rapidly activate in response to extracellular acidification. They are members of the ENaC/DEG superfamily and consist of six subtypes in humans (ASIC1a, ASIC1b, ASIC2a, ASIC2b, ASIC3 and ASIC4), with the widely expressed ASIC1a homotrimers being the most studied. ^7,8^

In neurons, activation of ASIC1a-mediated sodium currents contributes to electrical signals underlying a variety of brain functions, including long-term potentiation, which is important in learning and memory.^9,10^ In contrast, sustained ASIC1a activation under conditions of ischemia and metabolic acidosis is thought to contribute to neurotoxicity.^11^ Additionally, ASIC1a has gained attention in cancer research due to its ability to sense the acidic tumor microenvironment.^12^

ASIC1a has been well-characterized in heterologous expression systems, but critical gaps remain in our understanding of how they function in physiological and pathophysiological settings. For example, recombinant ASIC1a requires abrupt pH drops below 7.0 for activation, a threshold usually incompatible with normal fluctuations in tissue pH.^13^ Additionally, recombinant ASIC1a undergoes fast and complete desensitization within seconds of its activation.^8–11^ However, since the discovery of proton-mediated currents, it has been evident that proton sensitivity and current kinetics can vary significantly across different cell types and within cell populations.^2^ In fact, dorsal root ganglia, as well as different cancerous cells of the nervous system, such as neuroblastoma and glioblastoma cells, display ASIC-mediated currents that are characterized by slow and incomplete desensitization, along with increased proton sensitivity.^2,16–18^ Differences in ASIC subtype expression have been proposed to partially explain this heterogeneity across cell types^16,17,19^, but the evidence for this notion remains inconclusive and fails to fully account for the range of observed behaviours. We therefore hypothesize that at least part of this biologically relevant variability originates from cell type- or cell population-specific protein-protein interactions (PPIs).

Motivated by this hypothesis, we identify monocarboxylate transporter 1 (MCT1) as a subtype-specific modulator of ASIC1a. Strikingly, this ASIC1a-MCT1 coupling enhances proton sensitivity and gives rise to slow and incomplete desensitization of ASIC1a-mediated currents. Through domain-swapping between subtypes within the ASIC and MCT family members, we delineate the molecular mechanism of the functional ASIC1a-MCT1 coupling. Furthermore, we find that in a cancer stem cell line derived from human Glioblastoma multiforme (GBM), MCT1 is absolutely required for the atypical non-desensitizing proton-evoked currents. Remarkably, endogenous expression of ASIC1a together with MCT1 significantly prolongs the lifespan of mice intracranially injected with glioblastoma cancer stem cells, revealing a surprising transporter-ion channel interaction with potential implications in tumour biology. This discovery clarifies at least part of the molecular origin for the long-established functional diversity of ASIC1a in the nervous system, provides new insight into ASIC1a regulation by PPIs, as well as pointing towards potential therapeutic implications.^20,21^

## Results

### MCT1 enhances pH sensitivity and slows desensitization of ASIC1a

We hypothesized that atypical ASIC1a-mediated currents observed in different cell types and populations may be caused by yet unknown PPIs of ASIC1a. To test this hypothesis directly, we aimed to identify protein interaction partners of human ASIC1a in HEK293 cells, which display robust endogenous ASIC1a expression.^22^ We therefore expressed human ASIC1a, C-terminally fused to eGFP as a pull-down handle (ASIC1-GFP), in HEK293T cells lacking endogenous ASIC1a (HEK ASIC1^-/-^)^23^. The interactome of ASIC1a was then determined using a GFP nanobody for the immunoprecipitation of ASIC1a-GFP, followed by high-resolution proteomic analysis. The human monocarboxylate transporter 1 (MCT1) was consistently co-purified with ASIC1a-GFP in three independent experiments (Fig S1A). MCT1 is responsible for the bidirectional transport of monocarboxylates, such as L-lactate, utilizing protons as the driving force for transport. As such, it plays a central role in regulating cellular metabolism, as well as extracellular and intracellular pH.^24,25^ Notably, both MCT1 and ASIC1a rely on protons for transport and activation, respectively.

To assess if the ASIC1a-MCT1 interaction affects ASIC1a function, we co-expressed ASIC1a with MCT1 in HEK ASIC1^-/-^ cells and conducted whole-cell patch clamp recordings using a fast perfusion system. To compare pH_50_ values for ASIC1a in the absence and presence of MCT1 we obtained proton concentration-response curves (CRC) by applying recording solutions with different proton concentrations (Fig. 1A-B). We found that the proton CRC was significantly left-shifted in the presence of MCT1, demonstrating that ASIC1a was more sensitive to protons when co-expressed with MCT1 (pH_50_: 6.6 [6.61-6.66] and 6.9 [6.89-6.94] for ASIC1a alone and with MCT1, respectively, (p<0.0001, unpaired t-test)). This striking difference in proton sensitivity is best illustrated using the ratio of current amplitudes at pH 6.9 to pH 5.6 (Fig 1C and table S1), which we hence used as a proxy for pH sensitivity in all subsequent experiments. Besides proton sensitivity, the current responses recorded in the absence and presence of MCT1 revealed a drastic difference in the desensitization profile. At a saturating concentration of protons (pH 5.6), we found that MCT1 slowed ASIC1a desensitization kinetics significantly (K_des_: 0.72 ± 0.15 s^-1^ and 0.18 ± 0.06 s^-1^ for ASIC1a alone or with MCT1, respectively; p<0.0001, unpaired t-test) (Fig 1C-D and table S1). Furthermore, channel desensitization was incomplete in the presence of MCT1, resulting in a significant increase in steady-state current (SSC) during prolonged activation for 40 s (from 3 ± 5% to 44 ± 17% of peak current for ASIC1a alone and with MCT1, respectively; p<0.001, unpaired t-test) (Fig 1C-D). By contrast, MCT1 had no significant effect on the amplitudes of ASIC1a-mediated current responses (p=0.990, unpaired t-test) or on recovery from desensitization after 10 s or 30 s of wash-out in pH 8 (p=0.947, unpaired t-test). Furthermore, we found no differences between ASIC1a alone and in the presence of MCT1 on the current-voltage (IV) index recorded by measuring current amplitudes induced by pH 5.6 at -40mV and +40mV (p=0.420, unpaired t-test) (Fig S1B). These results show that the presence of MCT1 has a profound functional impact on ASIC1a and that this does not arise from altered channel abundance on the cell surface.

**Figure 1.**
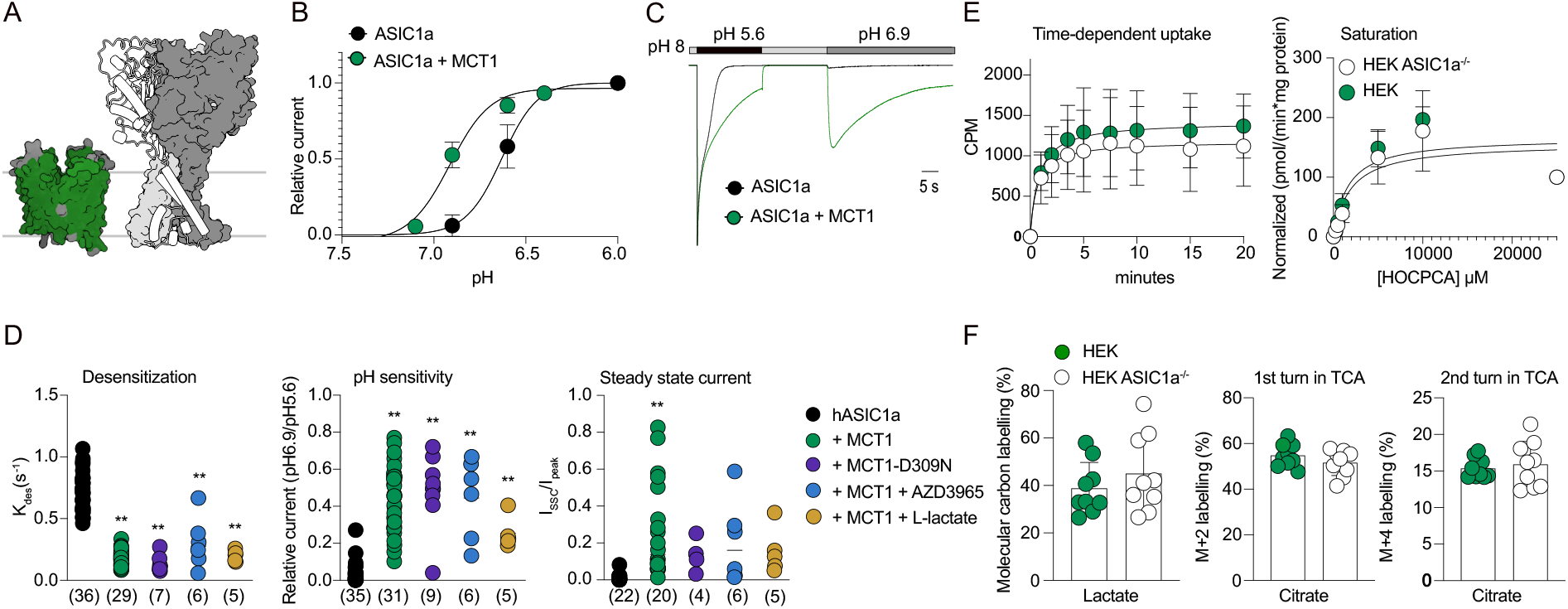
MCT1 affects ASIC1a function. **A** Structures of MCT1 (green) in outward open conformation (PDB # 6LZ0)^26^ and ASIC1a (grey) in resting conformation^30^. Each of the three ASIC1a protomers consist of an intracellular N-terminus (not structurally resolved), followed by a re-entrant helix, PreM1, connected to transmembrane helix M1. M1 is followed by a large extracellular domain (ECD), connected to transmembrane helix M2, and followed by the intracellular C-terminus (not structurally resolved). **B** ASIC1a proton concentration response curves (CRC) in absence (black circles) and presence of MCT1 (green circles) in HEK ASIC1a^-/-^ cells. The current amplitude at each concentration was normalised to the current amplitude at pH 5.6, with each concentration tested on minimum four individual patches. **C** Representative current traces of whole-cell patch clamp recordings from HEK ASIC1^-/-^ cells overexpressing ASIC1a alone (black trace) and with MCT1 (green trace). **D** ASIC1a functional modulation is independent of MCT1 conformation. Summary of desensitization kinetics (left), pH sensitivity (middle) and steady state current (SSC) (right) of ASIC1a alone (black circles), with MCT1 (green circles), with MCT1-D309N (purple circles) locked in inward open conformation, in presence of AZD3965 (50 μM) (blue circles), locking MCT1 in outward-open conformation, and in presence of L-lactate (5 mM) (yellow circles). Asterisks denote significance levels in comparison to ASIC1a alone: **p < 0.0001, unpaired t-test. Number of repetitions in brackets. **E** Uptake of [^3^H]HOCPCA via endogenous MCT1 at pH 6. Summary of three repetitions of [^3^H]HOCPCA uptake over time (left) and saturation assay (right) using HEK WT cells (green circles) and HEK ASIC1a^-/-^ cells (white circles), with error bars as standard deviation. **F** The presence (green circles) or absence of endogenous ASIC1a (white circles) had no effect on ^13^C-L-lactate uptake via endogenous MCT1 or on incorporation of ^13^C-carbon in TCA metabolites. Graphs showing normalised uptake of ^13^C-L-lactate to total amount of intracellular L-lactate or incorporation of ^13^C-carbon in TCA cycle metabolites, normalised to total amount of the metabolite, here illustrated as ^13^C-carbon labelling in citrate 1^st^ turn, entry point of TCA cycle, and citrate 2^nd^ turn in TCA cycle. Three replicates/condition repeated on three individual experiment days.

Next, we sought to test if and how different functional states of MCT1 could affect the modulation of ASIC1a function. Based on the published structures of MCT1, we locked MCT1 in an outward-open conformation using the MCT1-selective inhibitor AZD3965^26^ or in the inward-open conformation by neutralizing the proton-coupling site (e.g. MCT1-D309N)^26^. We found that MCT1 had the same dramatic effect on ASIC1a function regardless of its conformational state (Fig 1D, S2A and table S1). This suggests that the functional modulation of ASIC1a is induced by the physical interaction between MCT1 and ASIC1a.

### MCT1 modulation affects ASIC1a pharmacology

To further investigate the extent to which MCT1 influences ASIC1a, we examined the effects of two well-established ASIC1a modulators, amiloride and PcTx1, which act through distinct mechanisms.^20^ The pore blocker amiloride inhibited ASIC1a function in the absence and presence of MCT1 in a comparable manner (current inhibition by amiloride for ASIC1a alone and with MCT1 was 39 ± 22% and 35 ± 20%, respectively (p=0.772, unpaired t-test)) (Fig S2B). PcTx1, on the other hand, is a gating modifier that increases ASIC1a pH sensitivity, but when applied at neutral pH (pH 7.4) inhibits subsequent activation by inducing desensitization.^27^ In the presence of MCT1, PcTx1 significantly enhances the non-desensitizing ASIC1a currents at pH 7.4, relative to ASIC1a alone (current at pH 7.4 relative to pH 5.6 for ASIC1a alone and ASIC1a with MCT1 was 25 ±17% and 48 ± 8%, respectively (p<0.05, unpaired t-test)) (Fig S2C). Thus, PcTx1 also acts as a gating modifier of ASIC1a in the presence of MCT1 but already markedly activates the channel at physiological pH.

In addition, L-lactate, the major substrate for MCT1, has previously been shown to increase ASIC1a-mediated currents by chelating divalent ions that would otherwise inhibit the channel.^28^ While the concentration of L-lactate required for chelation is typically above physiological levels (15 mM), we aimed to investigate whether physiological concentrations of L-lactate (5 mM) could influence ASIC1a mediated currents when in the presence of MCT1. We found that L-lactate did not further affect ASIC1a desensitization or maximum current at pH 5.6 during whole-cell patch clamp recordings in presence of MCT1 (I_max_: 5.23 ± 3.95 nA and 4.67 ± 2.47 nA, (p=0.660, unpaired t-test), K_des_: 0.20 ± 0.04 and 0.18 ± 0.06 (p=0.635, unpaired t-test)) (Fig 1D & S2B). A small yet significant effect on pH sensitivity was found and attributed to variability between individual patch-clamp recordings (pH sensitivity measured as current at pH 6.9 normalised to current at pH 5.6 (in percentage): 25 ± 9% and 44 ± 17% for ASIC1a with MCT1 +/- L-lactate (5 mM), respectively (p<0.05, unpaired t-test)) (Fig 1D).

### ASIC1a has no major impact on MCT1 transport activity

As the presence of MCT1 affected ASIC1a function, we speculated if conversely ASIC1a would alter MCT1 activity. We therefore tested the uptake capacity of MCT1 in the absence and presence of ASIC1a by incubating wild-type (WT) HEK293T cells, i.e. cells endogenously expressing both MCT1 and ASIC1a, and HEK ASIC1a^-/-^ cells (see above) with the MCT1-selective radioligand [^3^H]HOCPCA. ^29^ Time-dependant uptake was linear up to approximate 5 minutes using a radioligand concentration of 100 nM with half-time to maximum at similar values (0.76 ± 0.22 min and 0.83 ± 0.17 min for HEK ASIC1^-/-^ and HEK WT, respectively (p=0.467, unpaired t-test)) (Fig 1E and S3A). Subsequent saturation experiments revealed no significant differences between absence and presence of ASIC1a (Km: 1.6 [0.81-3.06] mM and 1.87 [0.83-4.19] mM (p=0.4697, unpaired t-test), and Vmax: 166 [137-200] pmol/min*mg protein and 157 [123-198] pmol/min*mg protein for HEK WT and HEK ASIC1a^-/-^, respectively (p=0.127, unpaired t-test)) (Fig 1E). To test if ASIC1a would affect L-lactate uptake and following metabolism we incubated HEK WT and HEK ASIC1a^-/-^ cells with ^13^C-labelled L-lactate for 30 minutes at pH 6, followed by gas chromatography-mass spectrometry (GC-MS) analysis. We found no significant difference in ^13^C-L-lactate uptake (HEK WT: 39 ± 11%, HEK ASIC1a^-/-^: 45 ± 16%, p=0.361,unpaired t-test), or in incorporation of ^13^C-carbon in TCA cycle metabolites, such as citrate, α-ketoglutarate and alanine (Fig 1F and S3B). We therefore conclude that the presence of ASIC1a does not have a major functional impact on MCT1.

### MCT2 and MCT4 also modulate ASIC1a function

MCT1 belongs to a family of four proton-coupled monocarboxylate transporters in humans, MCT1 to MCT4, and we wondered if other MCT family members could modulate the function of ASIC1a. While MCT3 is only found in the retinal pigmented epithelium and the choroid plexus epithelium^25^, MCT1, MCT2 and MCT4 are widely expressed in the central and peripheral nervous system, as is ASIC1a.^25^ Therefore, in addition to MCT1, we also investigated if MCT2 and/or MCT4, predominantly expressed in neurons and astrocytes, respectively, would affect ASIC1a function. Whole-cell patch clamp recordings of HEK ASIC1a^-/-^ cells co-expressing ASIC1a and MCT2 gave rise to currents with slower desensitization (K_des_: 0.47 ± 0.09 s^-1^, at pH 5.6 (p<0.001, unpaired t-test)) and a small, but non-significant increase in pH sensitivity (p = 0.98, unpaired t-test) (Fig 2A and C). Contrary to MCT1 and MCT2, MCT4 slightly increased ASIC1a current desensitization (0.80 ± 0.24 s^-1^, at pH 5.6, p=0.96,unpaired t-test) and slightly decreased sensitivity to pH 6.9 relative to ASIC1a alone (p = 0.96, unpaired t-test), thus exhibiting a trend towards inverse functional effects compared to MCT1/2.

**Figure 2.**
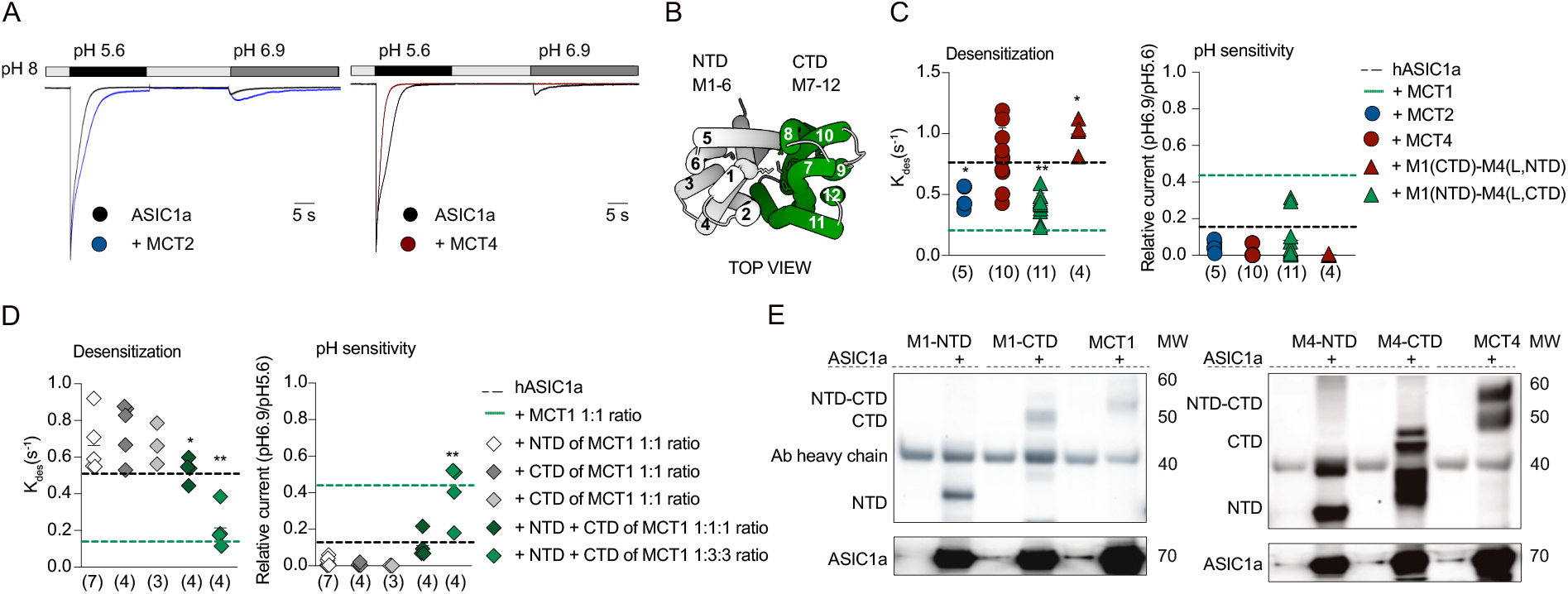
MCT-CTD dictates modulatory effect on ASIC1a. **A** Representative current traces of ASIC1a alone (black trace) and in the presence of MCT2 (blue) or MCT4 (red) in response to application of pH 5.6 and 6.9. **B**. Top view of MCT1 (PDB # 6LZ0), color-coded to highlight the NTD (grey) and CTD (green), with numbered transmembrane helices. **C** Summary of desensitization kinetics (left) and pH sensitivity (right) of ASIC1a alone (average shown as black dashed line), ASIC1a with MCT1 (average shown as green dashed line), with MCT2 (blue circles), with MCT4 (red circles), M4-NTD-M1-CTD (green triangles) and M1-NTD-M4-CTD (red triangles). Asterisks denote significance levels in comparison to ASIC1a alone: *p < 0.05 and **p < 0.0001, unpaired t-test. Number of repetitions in brackets. **D** CTD of MCT1 alone does not modulate ASIC1a function but is dependent on the presence of NTD. Dot plots summarizing the effect on desensitization kinetics at pH 5.6 (left) and pH sensitivity (right) of ASIC1a alone (average illustrated by black dashed line), with MCT1 (average illustrated by green dashed line), for ASIC1a co-expressed in 1:1 ratio with the MCT1-NTD (white diamonds), MCT1-CTD (dark grey diamonds) and MCT1-NTD + CTD (dark green diamonds) or in 1:3 ratio of ASIC1a with MCT1-CTD (grey diamonds) and MCT1-NTD + CTD (green diamonds). A 1:3 ratio (measured in µg DNA) of ASIC1a with MCT1-NTD + MCT1-CTD is necessary to rescue the full effect elicited by full-length MCT1. Asterisks denote significance levels in comparison to ASIC1a alone: *p < 0.05 and **p < 0.0001, unpaired t-test. Number of repetitions in brackets. **E** Western Blot illustrating the pull down of MCT1 (left) and MCT4 (right) as well as their individual truncated versions comprised of only NTD or CTD of MCT1/MCT4 using ASIC1a as bait. Pull-down results indicate a broader interaction surface on MCTs. Immunoprecipitation followed by western blotting was repeated three times.

### The MCT C-terminal domain dictates modulation of ASIC1a function

The divergent effects of MCT1 and MCT4 on ASIC1a function prompted us to investigate the structural determinants that dictate the type of functional modulation on ASIC1a. MCT1 and MCT4 share only 39.4% sequence identity but have a common topology consisting of 12 transmembrane helices. The first six transmembrane helices form the N-terminal domain (NTD), while the latter six form the C-terminal domain (CTD), connected by a large intracellular loop (Fig 2B).^24,26^ Thus, chimeric constructs comprising the NTD (M1-M6) of either MCT1 or MCT4, paired with the CTD (M7-M12) from the other subtype were expressed together with ASIC1a in HEK ASIC1a^-/-^ cells (Fig 2B). We then conducted whole-cell patch clamp recordings to test for changes in current desensitization and/or pH sensitivity. Notably, the chimeric construct incorporating the CTD of MCT1 (M1(CTD)-M4(L,NTD) exhibited MCT1-like effects, while the construct containing the CTD of MCT4 (M1(NTD)-M4(L,CTD) displayed MCT4-like effects on ASIC1a desensitization kinetics and pH sensitivity (Fig 2C and table S1). Thus, the CTD of the transporter dictates the nature of the modulatory effect on ASIC1a. Additional chimeric constructs of MCT1 with the intracellular C-terminus of MCT4 (M1(1-12)-M4(C-terminus)), or MCT1 with the intracellular M6-M7 linker of MCT4 (M1(1-12)-M4(L)), both showed MCT1-like effects on ASIC1a function, confirming that the modulation is caused by the transmembrane helices of the MCT-CTD (Fig S4B and table S1). However, co-expressing ASIC1a with the CTD of MCT1 alone did not affect ASIC1a currents. We only observed the full MCT1 effect on ASIC1a when co-expressing both the CTD and NTD of MCT1 *in trans* (Fig 2D and table S1). In addition, we found that both MCT1 and MCT4, as well as the truncated domains of only the NTD or CTD of MCT1 and MCT4, all co-immunoprecipitated with ASIC1a, indicating a broad interaction surface covering transmembrane helices on both the NTD and CTD (Fig 2E). Therefore, and despite the common topologies of the MCT family, we have uncovered diverging functional effects on ASIC1a mediated by the transporter CTD.

### MCT1 functionally modulates ASIC1a subtype selectivity via PreM1

The question whether MCT1 can modulate other ASIC subtypes is crucial to uncover potential broader regulatory mechanisms. Thus, we co-expressed MCT1 with either ASIC1b, ASIC2a or ASIC3 (sharing between 46.8 and 72.5% sequence identity with ASIC1a) in HEK ASIC1a^-/-^ cells and conducted whole-cell patch clamp recordings and immunoprecipitation assays. We found that neither ASIC1b, nor ASIC2a or ASIC3 can be functionally modulated by MCT1 (Fig 3A and S5A-B). However, co-immunoprecipitation experiments for MCT1 with ASIC2a and ASIC3 indicated that MCT1 likely interacts with both proteins (Fig S5C). Since the modulation induced by MCT1 is highly selective for the ASIC1a subtype, we speculated whether human MCT1 could influence ASIC1a from different species. Thus, we co-expressed human MCT1 with mouse and chicken ASIC1a, respectively, and conducted whole-cell patch-clamp recordings. Notably, human ASIC1a shares 97.9% sequence identity with mouse ASIC1a but only 63.8% with chicken ASIC1a. Consistently, functional modulation by human MCT1 was only observed with mouse ASIC1a (Fig S5D-E).

**Figure 3.**
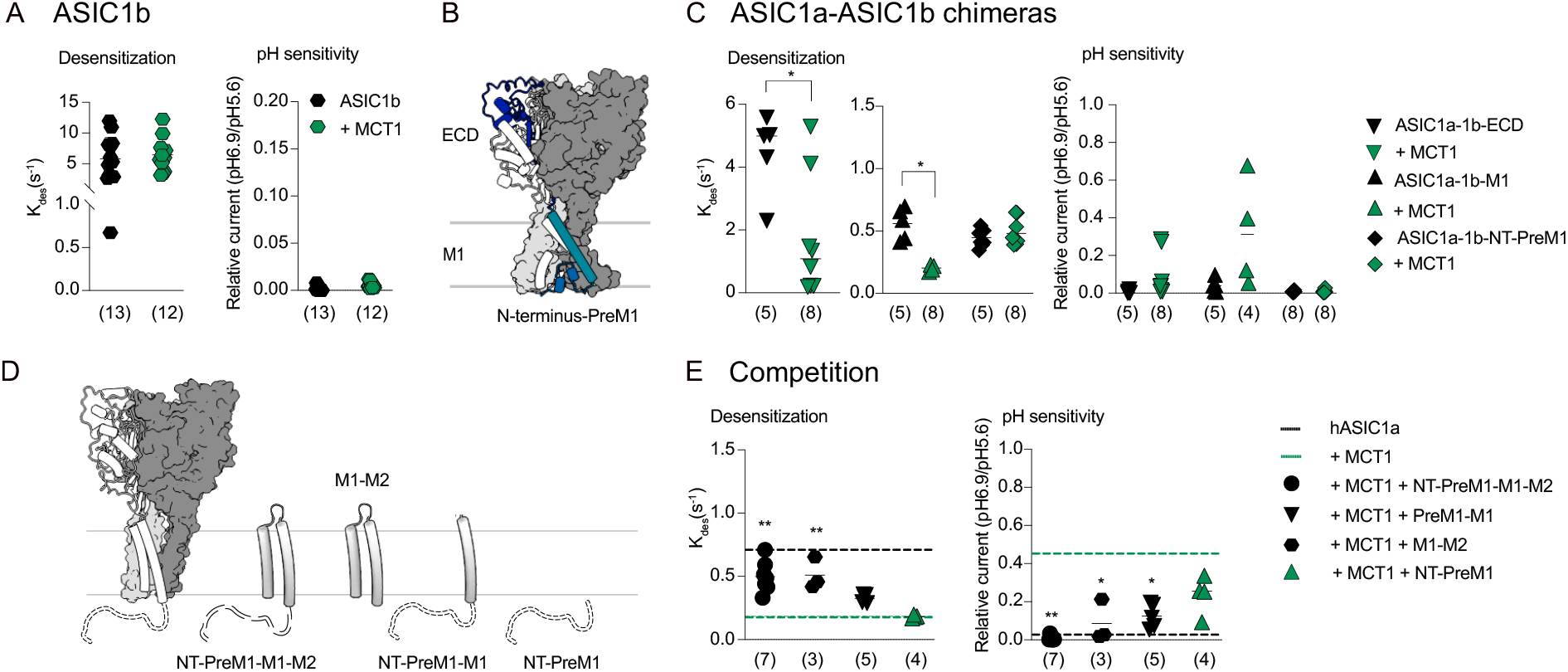
MCT1 selectively modulates ASIC1a function via PreM1. **A** ASIC1b is not affected by MCT1. Summary of desensitization kinetics at pH 5.6 (left) and pH sensitivity (right) of hASIC1b alone (black hexagons) and co-expressed with MCT1 (green hexagons). **B** Structure of ASIC1a in resting conformation. Non-conserved elements between ASIC1a and ASIC1b are highlighted in shades of blue; PreM1 (blue), M1 (light blue) and the first part of ECD (dark blue). **C** Complete loss of MCT1 modulation only occurs for ASIC1a chimeric construct containing the N-terminus-PreM1 of ASIC1b (ASIC1a-1b-NT-PreM1). Summary of desensitization kinetics at pH 5.6 (left) and pH sensitivity (right) of ASIC1a-ASIC1b chimeric constructs alone (black) and with MCT1 (green); ASIC1a with ASIC1b N-terminus-PreM1 (triangles), ASIC1a with ASIC1b M1 (diamonds), ASIC1a with ASIC1b ECD (opposite triangles). Asterisks denote significance levels in comparison to ASIC1a-ASIC1b chimeric construct alone: *p < 0.05, unpaired t-test. Number of repetitions in brackets. **D-E** Outcompeting the interaction between ASIC1a and MCT1 using mini-proteins consisting of TMD segments of ASIC1a indicates that M1 of ASIC1a is involved in the coupling with MCT1. Illustration of miniproteins of ASIC1a segments (**D**). Summary of desensitization kinetics at pH 5.6 (left) and pH sensitivity (right) (**E**) for ASIC1a when co-expressed with MCT1 and the following TMD segments of hASIC1a; PreM1-M1-M2 (black circles), PreM1-M1 (back hexagons) or M1-M2 (black triangles) and N-terminus-PreM1 (green triangles). Asterisks denote significance levels compared to ASIC1a with MCT1 alone: * p < 0.05 and **p < 0.05, unpaired t-test. Number of repetitions in brackets.

Each ASIC1a protomer consists of intracellular N- and C-termini, a large extracellular ligand-binding domain (ECD), and two transmembrane helixes (M1 and M2), that form the ion-conducting pore. In the resting state, part of the N-terminus (from AA15-AA40) of ASIC1a can adopt a membrane embedded re-entrant loop, termed PreM1, that packs into the ion channel pore and likely interacts with pore-lining residues in M2.^30–32^ Upon binding of protons to the ECD, the channel opens, likely allowing the PreM1 to unwind and extrude into the intracellular environment.^30^ ASIC1a and ASIC1b share 72.5 % sequence identity, with only the N-terminus, PreM1, M1 and the proximal part of the ECD displaying sequence differences (Fig 3B). Because ASIC1b was not affected by MCT1 (Fig 3A), this presented a unique opportunity to explore the molecular basis subtype-specific functional modulation of ASIC1a by MCT1. We therefore designed three chimeric constructs by swapping segments between ASIC1a and ASIC1b. The first construct comprised ASIC1a with the ECD of ASIC1b (ASIC1a-1b-ECD), the second featured ASIC1a with transmembrane helix M1 of ASIC1b (ASIC1a-1b-M1), and the third consisted of ASIC1a with the N-terminus and PreM1 of ASIC1b (ASIC1a-1b-NT-PreM1) (Fig 3B). Each construct was then expressed alone or together with MCT1 in HEK ASIC1^-/-^ cells for whole-cell patch clamp recordings. We found that only ASIC1a containing the N-terminus and PreM1 of ASIC1b (ASIC1-1b-PreM1) could no longer be modulated by MCT1 (K_des:_ 0.45 ± 0.07 s^-1^ and 0.48 ± 0.11 s^-1^ for ASIC1a-1b chimera alone and with MCT1, respectively (p=0.498, unpaired t-test)) (Fig 3C and table S1). This underlined the importance of the N-terminus-PreM1 in the subtype-selective modulation by MCT1. Additionally, to rule out a contribution by the C-terminus, we examined an ASIC1a construct with the intracellular C-terminus truncated at position 466 (ASIC1aΔ466). When overexpressed in HEK ASIC1^-/-^ cells, MCT1 was still able to functionally modulate ASIC1aΔ466 in whole-cell patch-clamp recordings (Fig S5F and table S1). This shows that, unlike the N-terminus-PreM1 region, the C-terminus does not play a role in ASIC1a coupling.

Thus, we predict that the interaction surface on ASIC1a could be either the N-terminus-PreM1 region or the M1 and/or M2 helices located near the N-terminus-PreM1 of ASIC1a. We thus sought to design miniproteins comprising small parts of the ASIC1a TMD region to evaluate their capacity to compete for interaction with MCT1. Four miniproteins, N-terminus-PreM1 (NT-PreM1), N-terminus-PreM1-M1 (NT-PreM1-M1), M1 connected to M2 via short linker (M1-M2), and N-terminus-PreM1-M1 linked to M2 (NT-PreM1-M1-M2) of ASIC1a, were individually co-expressed with ASIC1a and MCT1 in HEK ASIC1a^-/-^ cells, followed by whole-cell patch-clamp recordings (Fig 3D-E). All miniproteins containing the M1 helix of ASIC1a could eliminate the effect of MCT1 on ASIC1a function, albeit to varying degrees, indicating the potential involvement of M1 of ASIC1a in the ASIC1a-MCT1 interaction (Fig 3D-E).

### Knocking out MCT1 in human glioblastoma affects proton induced currents

A subset of cancer cells from Glioblastoma multiforme (GBM), the most lethal brain cancer in adults, exhibit atypical proton-mediated currents characterized by slow and incomplete desensitization, along with increased proton sensitivity.^16–18^ To determine whether ASIC1a-MCT1 coupling might be the cause for the atypical currents observed in glioblastoma cells we sought to determine the relative abundance of ASIC subtypes and MCT family members in the Glioblastoma map (GBmap) of single-cell RNA sequencing data^33^. We found that in malignant brain cells, ASIC1a and MCT1 are the dominant subtypes among ASIC and MCT family members, respectively (Fig S6A). In addition, we found that only 23% of ASIC1a-expressing healthy glial-neuronal cells co-express MCT1, while 47% of ASIC1a-expressing malignant cells co-express MCT1 (Fig S6B). With this in mind, we selected the R54 glioblastoma cancer stem cell line, which displays proneural expression pattern and represents one the most reliable in vitro model currently available for studying GBM biology.^16,34,35^ In line with previously published data^16^, whole-cell patch-clamp recordings of R54 cells gave rise to three distinct types of proton-induced currents. Type I showed a peak current that rapidly and completely desensitized at pH 5.6 (K_des_: 1.16 ± 0.41 s^-1^, SSC: 0.4% of peak current) and no or negligible currents at pH 6.9 (Fig 4A and B), thus resembling proton-induced currents from HEK293T cells overexpressing ASIC1a alone. Type II currents showed fast but incomplete desensitization, giving rise to SSC during prolonged activation at pH 5.6 (K_des_: 0.97 ± 0.34 s^-^1, SSC: 23% of peak current). Activation by pH 6.9 resulted in inward currents with amplitudes of approximate 25 % of its full activation by pH 5.6. (Fig 4A and B). Type III currents showed minimal desensitization, followed by large steady-state currents (K_des_: 1.14 ± 0.34 s^-^1, SSC: 56% of peak current), resulting in almost square currents at pH 5.6. The currents induced by pH 6.9 resulted in inward currents with amplitudes equal to or twice as large as currents induced by pH 5.6 (Fig 4A and B). The relative occurrence of the different current types in WT R54 cells were 57% for type I, 32% type II and 11 % type III, with type II and type III currents showing a striking resemblance to proton-induced currents from HEK cells overexpressing ASIC1a together with MCT1. We therefore hypothesized that deletion of MCT1 in R54 cells would decrease the number of cells giving rise to the atypical type II and III proton induced currents. To test this notion directly, we used CRISPR/Cas to generate MCT1 KO R54 cells (Fig S6). Remarkably, we found that the number of cells exhibiting type I currents drastically increased from 57% to 96 % in absence of MCT1, whereas only 4% showed type II currents and none of the cells gave rise to type III currents (Fig 4C). These results strongly indicate that ASIC1a-MCT1 coupling gives rise to the atypical proton-induced currents observed in WT R54 cells.

**Figure 4.**
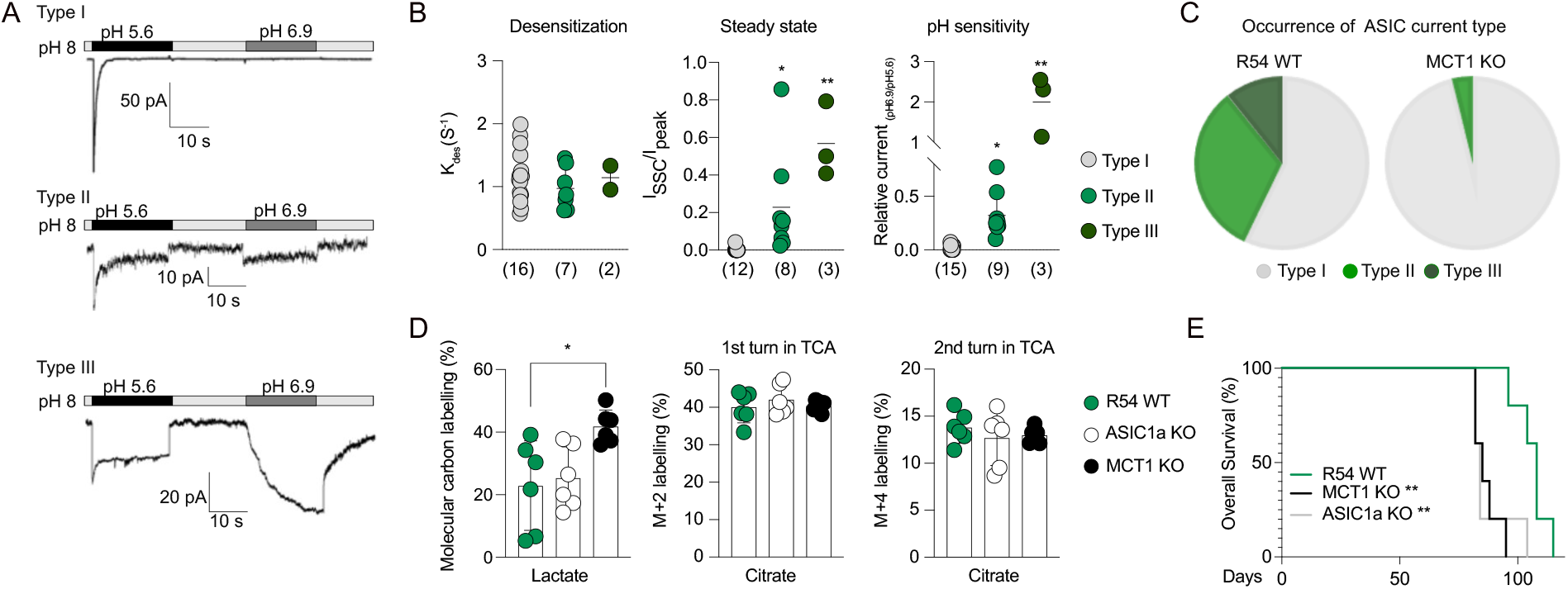
MCT1 modulates ASIC1a function in subpopulation of human glioblastoma. **A** Traces illustrating three types of currents recorded by applying pH 5.6 and pH 6.9 to R54 cancer stem cells. Type I currents (upper trace) were similar to currents from HEK cells overexpressing ASIC1a. Type II currents (middle trace) show incomplete desensitization, resulting in increased steady state currents upon prolonged exposure to pH 5.6 and increased pH sensitivity, similar to the observed change on ASIC1a function when co-expressed with MCT1 in HEK cells. Type III (lower trace) showing little to no desensitization, resulting in square currents at pH 5.6 and increased pH sensitivity with current amplitudes at pH 6.9 equal to or bigger than currents induced by pH 5.6. **B** Summary of desensitization kinetics (left), steady state currents (middle) and pH sensitivity (right) for type I (grey), type II (green), type III (dark green) currents. Asterisks denote significance levels compared to type I currents: * p < 0.05 and **p < 0.001, unpaired t-test. Number of repetitions in brackets. **C** Pie charts showing the relative occurrence of current types recorded from 28 R54 WT cells (left) and 27 MCT1 KO R54 cells (right). **D** Bar graphs showing uptake of ^13^C-labelled L-lactate via endogenous MCTs in R54 cells at pH 6 and following incorporation of ^13^C-labelled carbon in TCA cycle metabolites, illustrated as ^13^C-labelled citrate 1^st^ round (entry point of TCA cycle) and citrate 2^nd^ round (one turn in the TCA cycle). ASIC1a KO R54 (white circles) showed similar uptake and subsequent incorporation of ^13^C-labelled carbon in TCA metabolites as WT R54 (green circles). In contrast, MCT1 KO R54 (black circles) showed an increase in uptake of ^13^C-labelled L-lactate (p>*0.05, unpaired t-test) which was not detected as incorporation into TCA metabolites. Two replicates/condition was repeated on three individual experiment days. **E** Kaplan-Meier curves showing days of survival of five mice injected with WT R54 (black), five mice with ASIC1a KO R54 (grey) and five mice with MCT1 KO R54 (black). The mice injected with ASIC1a KO R54 and MCT1 KO R54 died significantly faster than mice injected with WT R54 cells (** p<0.01, Mantel-Cox Test).

### ASIC1a has no impact on lactate uptake and metabolism in glioblastoma

Cancer cells are characterized by high metabolic rates, and it has recently been suggested that targeting the proton-assisted L-lactate uptake via MCT1 or lactate metabolism in glioblastoma could be a novel therapeutic approach.^36,37^ In R54 cells, endogenous MCT1 plays a significant role in modulating the currents from endogenous ASIC1a (Fig 4A-C). Therefore, we speculated if endogenous ASIC1a could affect MCT1-induced lactate uptake and metabolism in the R54 glioblastoma cells. Thus, L-lactate uptake and metabolism were assessed in WT R54 cells, MCT1 KO R54 cells, and ASIC1a KO R54 cells^38^ by incubating the cells with ^13^C-labeled L-lactate for 30 minutes at pH 6, followed by GC-MS analysis. We found no difference in uptake between WT R54 cells and ASIC1a KO R54 cells (23±14% and 25 ±10%, respectively (p=0.735, unpaired t-test)), indicating that ASIC1a does not affect L-lactate uptake or metabolism in R54 cells (Fig 4D). To our surprise, we found that MCT1 KO R54 showed a significant increase in L-lactate uptake (42 ± 5%) relative to uptake by WT R54 cells (p<0.05, unpaired t-test) (Fig 4D). However, the increase in L-lactate uptake was not translated into an increase in ^13^C-labelled carbon in TCA cycle metabolites (Fig 4D and S6D-E), suggesting that lactate accumulates in MCT1 KO R54 cells. We conclude that ASIC1a, or its coupling with MCT1, does not contribute to proton-assisted L-lactate uptake or metabolism in R54 cells. In contrast, absence of MCT1 in R54 cells enhances L-lactate uptake, which may serve as an epigenetic metabolite driving pro-survival gene expression. This paradoxical effect of MCT1 silencing represents an important consideration when targeting GBM.

### ASIC1a enhances survival in mouse xenograft model

Acidic tumour microenvironment and metabolic reprogramming are hallmarks of cancer and both ASIC1a and MCT1 have been associated with cancer progression.^39,40^ However, in the specific case of glioblastoma, research has indicated that high ASIC1a expression is linked to prolonged lifespan. ^41,42^ Thus, we aimed to investigate how the presence or absence of the proton-sensitive ASIC1a and MCT1 in R54 cells would influence mouse survival after xenograft implantation in the brain. To this end, we assessed the lifespan of 15 mice intracranially injected with WT R54, ASIC1a KO R54, or MCT1 KO R54 cells. Given the aggressive nature of human glioblastoma cell lines in mice intracranial xenografts, we did not expect large survival differences between groups. Typically, untreated mice survive 30–90 days post-implantation, while treated mice can survive more than 100 days, depending on the treatment.^43,44^ We found that mice injected with ASIC1a KO cells died significantly faster (lifespan: 82-104 days) than mice injected with WT R54 cells (lifespan: 96-115 days) (Fig 4E), supporting previous findings indicating that the presence of ASIC1a is associated with prolonged overall survival in glioblastoma patients.^41,42^ Similarly, mice injected with MCT1 KO R54 cells died significantly faster (lifespan: 82-95 days) than mice injected with WT R54 cells, suggesting that the increase in intracellular L-lactate observed for MCT1 KO R54 cells, and perhaps the concomitant loss of ASIC1a modulation, indeed leads to increased mortality.

## Discussion

The origin of atypical endogenous ASIC-mediated currents observed in e.g. dorsal root ganglia and glioblastoma cells, and the conundrum of how ASIC channels can be rendered sufficiently proton-sensitive to become activated in e.g. synaptic environments that undergo less pronounced changes in pH have been long-standing questions of potentially high (patho)physiological relevance. A variety of PPIs have been suggested for different ASIC subtypes.^45^ However, the identification of the ASIC1-MCT1 interaction is particularly relevant as it is associated with a drastic increase in proton sensitivity, along with slow and incomplete desensitization of ASIC1a (Fig 1B-D). Our data not only demonstrate a coupling between ASIC1a and MCT1, but also show its involvement in generating atypical currents in glioblastoma cancer stem cells, and its contribution to enhanced survival in mouse xenograft models (Fig. 4A–C).

Acidic tumour microenvironments are common in cancer, with pH levels typically ranging from pH 6.2 to 7.1, depending on tumour aggressiveness and differences between the tumour core and periphery.^46–48^ Previous studies have reported conflicting results regarding the role of ASIC1a in cancer, some suggesting that increased ASIC1a expression is associated with metastasis and proliferation.^40,49^ In the case of glioblastoma, however, studies have shown that high ASIC1a expression is correlated with lifespan prolongation.^38,41,42^ In our work, we found that the presence of ASIC1a and MCT1 together significantly extends the lifespan of mice subjected to glioblastoma xenografts (Fig 4E). Various ASIC1a-induced pathways have been proposed to lead to suppressed glioblastoma cell growth, proliferation and invasion, accompanied by an increase in apoptosis and necroptosis.^38,40^ We suggest that increase in ASIC1a pH sensitivity and incomplete desensitization, caused by coupling with MCT1, contributes to glioblastoma cell death.

Metabolic reprogramming is a key characteristic of cancer and, consistent with previous studies^37,39,50^, our findings indicate that MCT1 expression is elevated in glioblastoma compared to non-malignant cells (Fig. S6A-B). In glioblastoma, MCT1 upregulation is associated with enhanced L-lactate uptake, as it serves as a cancer-promoting metabolite.^36^ Intriguingly, however, the limited clinical effect of the MCT1-specific inhibitor AZD3965 in cancer treatment has been partly attributed to the compensatory upregulation of other MCT family members.^51,52^ Indeed, we found that absence of MCT1 in R54 cells (MCT1 KO R54) led to an increase in L-lactate uptake (Fig 4D). We therefore speculate that this could be caused by compensatory upregulation of other MCTs, likely MCT2, which has a higher affinity for L-lactate. Accumulation of L-lactate as a cancer-promoting metabolite, along with the concomitant loss of ASIC1a modulation, thereby explain the increase in mortality of mice after MCT1 KO R54 cell xenografts (Fig 4E).

Recombinantly expressed ASIC1a undergoes fast desensitization and even a modest drop in pH will result in a steady state-desensitized state that renders the channel unresponsive to subsequent activation by protons.^8,13,14^ These characteristics would limit sustained ASIC1a activity during prolonged tissue acidosis *in vivo*. Potentiation of ASIC1a activity has been partly attributed to e.g. the release of arachidonic acid, spermines and L-lactate during pathological conditions such as ischemia, thereby contributing to cell death.^19^ In this context, our finding that direct functional modulation of ASIC1a by MCT1 leads to sustained currents even during small drops in pH may help explain, at least in some instances, how ASIC1a can contribute to neuropathologies during prolonged tissue acidosis.

Notably, a recent publication suggested a potential interplay between ASIC1a and MCT2 in primary neuronal cultures.^53^ The authors found that uptake of L-lactate by MCT2 gave rise to intracellular acidification, triggering a localized extracellular pH drop due to proton extrusion via the Na^+^/H^+^ exchanger 1 (NHE1) from nearby astrocytes. This pH drop was suggested to be sufficient to activate ASIC1a when localized in close proximity to MCT2 in neuronal membranes.^53^ Our work shows that ASIC1a and MCTs indeed directly couple (Fig. 2C). The presence of 5 mM L-lactate had no additional impact on ASIC1a current potentiation in the presence of MCT1 in whole-cell patch clamp recordings (Fig. 1D and S2A). However, it is intriguing that the coupling between ASIC1a and MCT2 (and/or MCT1) not only leads to functional modulation of ASIC1a, but also promotes its activation *in vivo* by facilitating the uptake of L-lactate. In this context, we found that PcTx1, typically considered a neuroprotectant due to its ability to induce ASIC1a desensitization at pH 7.4, has the ability to activate ASIC1a at physiological pH 7.4, when ASIC1a is co-expressed with MCT1 (Fig S2C). Together, these findings imply that the specific cellular context adds complexity to the role of ASIC1a in neurons during ischemic or metabolic acidosis and provides a striking example of how PPIs can affect ASIC1a function and pharmacology.

As for the required molecular determinants on ASIC1a, our work suggests that modulation by MCT1 is highly ASIC1a-subtype selective, and that this originates from the ASIC1a N-terminus-PreM1 (Fig 3A-D). Several structural studies have shown that the ASIC1a TMD can exist in various different conformations. The PreM1 region has been structurally resolved in ASIC1a structures only when it forms a re-entrant helix within the ion channel pore, where it likely interacts with pore-lining residues in M2.^30,32^ Notably, mutations in the PreM1 region, e.g. T26V, result in ASIC1a currents similar to the currents observed for ASIC1a in presence of MCT1.^30,31^ Recent cryo-EM structures of human ASIC1a-T26V at pH 8.5 revealed that this mutation prevents the PreM1 from integrating into the channel pore, which might explain why the channel becomes more prone to opening even at proton concentrations that would not normally activate ASIC1a (pH 7), as well as slowing the desensitization.^30^ Similarly, we propose that MCT1 interferes with the ability of PreM1 to recoil into the channel pore, either through transient interactions with N-terminus-PreM1 outside the membrane region or by the direct physical contact with M1 of ASIC1a. Collectively, these findings support the notion that PreM1 can act as both an auto-modulatory domain, as well as engage in direct or indirect PPIs to alter for ASIC1a function.

In summary, we identify MCT1, MCT2, and MCT4 as novel interaction partners of ASIC1a that regulate its activity in different neuronal settings. The coupling between the MCT family and ASIC1a provides new insight into cell type-specific modulation of ASIC1a function under both physiological and pathophysiological conditions. In glioblastoma, where MCT1 expression is upregulated, we propose that atypical proton-induced ASIC1a currents contribute to promotion of cell death (Fig 5A). Disrupting this coupling via miniproteins or knocking out ASIC1a likely contribute to cancer progression due to the loss of ASIC1a-MCT1 mediated atypical currents (Fig 5B). Further, the absence of MCT1 in glioblastoma cancer stem cells leads to increased intracellular levels of L-lactate, a metabolite that promotes cancer growth, potentially via MCT2 with minimal impact on ASIC1a mediated currents. Collectively, this would contribute to the advancement of cancer progression (Fig 5C). This study represents a significant advance in understanding how cell type- and ASIC1a subtype-selective PPIs influence ASIC1a physiology and role in cancer biology.

**Figure 5.**
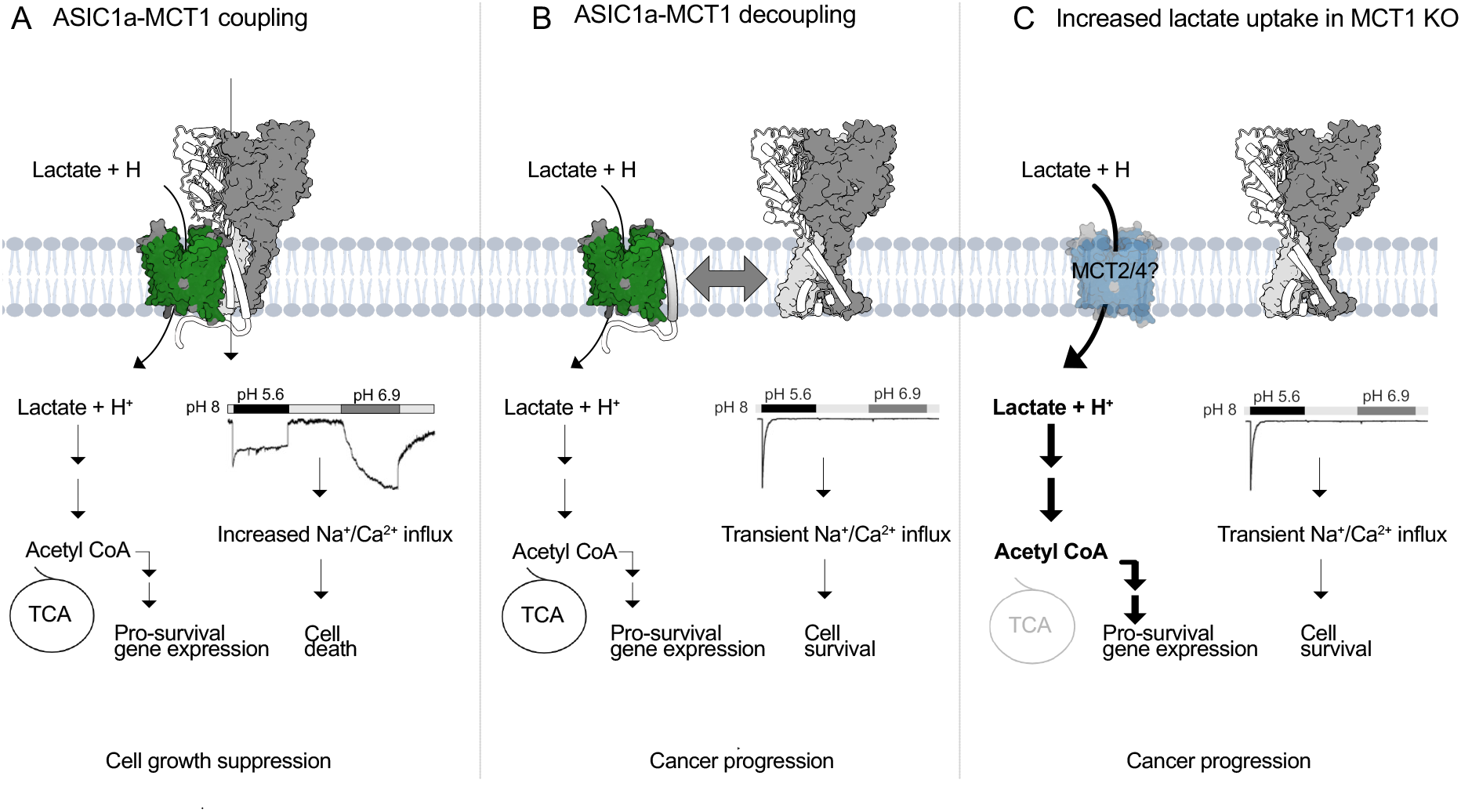
Proposed consequences of ASIC1a-MCT1 (de)coupling in glioblastoma. **A** Approximately 50% of R54 glioblastoma cancer stem cells (estimated by measuring occurrence of current types I-III) show ASIC-mediated currents with enhanced proton sensitivity, and slow and incomplete desensitization. These atypical proton-induced currents arise from the ASIC1a-MCT1 coupling, likely resulting in elevated intracellular Na^+^ and Ca^2+^ levels that promote cell death. Consequently, the ASIC1a-MCT1 interaction in R54 glioblastoma cancer stem cells slows cancer progression and may explain why high ASIC1a expression appears to enhance survival in glioblastoma patients and mouse xenograft models. **B** In the absence of ASIC1a (ASIC1a KO R54) or when the ASIC1a-MCT1 interaction is disrupted using miniproteins targeting the interaction surface on MCT1, fewer cells undergo cell death due to the lack of atypical proton-induced currents. This potentially explains why mice injected with ASIC1a KO R54 exhibit significantly faster mortality compared to those injected with R54 WT. **C** In the absence of MCT1 (MCT1 KO R54), L-lactate uptake is elevated. R54 cancer stem cells adapt to MCT1 loss potentially by compensating through the upregulation of other MCTs, such as MCT2, which has a higher affinity for L-lactate, and only minor effect on ASIC1a function. The rise in intracellular L-lactate likely accelerates tumour progression, potentially explaining the significantly increased mortality observed in mice injected with MCT1 KO R54 compared to those injected with R54 wild-type glioblastoma.

## Acknowledgements

We thank Janne Colding (Pless lab) for her outstanding technical assistance with molecular biology and cell culture work. We are grateful to Stefan Gründer for valuable input and for providing the R54 WT and R54 ASIC1a KO cells. We also acknowledge Javier Martin Gonzalez, manager of the Transgenic core facility at University of Copenhagen, for producing MCT1 KO R54 cells using CRISPR/Cas. Special thanks go to Dr Blanca Aldana, Dr Aisha Ameen and Heidi Nielsen for their input on L-lactate metabolism mapping and support in running samples for the GC-MS.

## Author contributions

*Conceptualization and Design*: Pless SA and Poulsen MH conceived the study and designed the experiments; *Data Collection*: Poulsen MH performed immunoprecipitation on GFP tagged ion channels, while Maurya S performed the MS and analysis. Poulsen MH, Ritter N and Heusser SA coducted whole-cell patch-clamp recordings on HEK cells, incl analysis. Poulsen MH, Kickinger S and Fisker-Andersen J conducted the HOCPCA radioligand uptake experiments and data analysis. Poulsen MH conducted the ^13^C-L-lactate uptake experiments on HEK and R54 glioblastoma cells. Chua HC conducted molecular biology. Usher S conducted analysis based on the single cell RNA sequencing atlas. Xue F performed co-immunoprecipitation expermients on ASICs and MCTs. Poulsen MH conducted immunoprecipitation experiments on R54 glioblastoma cells. Poulsen MH conducted whole-cell patch-clamp recordings on R54 glioblastoma cells. Haider S performed intracranial injections of R54 glioblastoma in mice and analysed the outcome; *Writing*: Poulsen MH drafted the manuscript, while all authors contributed with draft reviewing and editing; *Supervision*: Pless SA provided oversight, idea generation and mentorship throughout the project. The mass spectrometry was performed in the laboratory of Lundby A under supervision and guidance from Lundby A. HOCPCA uptake experiments was perfomed in the laboratory of Wellendorph P under supervision and guidance from Wellendroph P. The mice study was performed in the laboratory of Schramek D under supervision and guidance from Schramek D.

## Methods

### Molecular biology

The cDNA encoding all ASIC subtypes, human MCT subtypes, human P2×7 and human serotonin transporter (SERT), as well as chimeric constructs and channels fused with GFP or mCherry were acquired in pcDNA3.1+ from [TWIST Bioscience](http://www.twistbioscience.com/?utm_medium=cpc&utm_source=google&utm_campaign=PSR-GLBL-FY21-1808-MULTI-Branded&utm_term=twist%20bioscience&utm_content=kwd-301007611611) (see table S2 and S3 for construct information). DNA coding for ion channels was C-terminally tagged with a GFP, followed by 1D4 tag or incorporated into pcDNA3.1+ containing IRES GFP downstream of the channel for visualization in patch-clamp recordings and for protein purification in co-immunoprecipitation experiments. DNA coding for MCT constructs had an N-terminal mCherry and a Flag tag or incorporated into pcDNA3.1+ containing mCherry P2A upstream of the channel for visualization during patch-clamp recordings and protein purification in co-immunoprecipitation experiments. Site-directed mutagenesis was performed using PfuUltraII Fusion polymerase (Agilent) and custom DNA mutagenesis primers (Eurofins Genomics). All sequences were confirmed by sequencing of the full coding frame (Eurofins Genomics or Macrogen). For RNA transcription, cDNAs were linearized with XbaI, and capped cRNA was transcribed with the Ambion mMESSAGE mMACHINE T7 kit (Thermo Fisher Scientific).

### Cell lines and culturing

#### Human embryonic kidney cells

HEK293T cells were purchased from ATCC (Originally Cat# CRL-3216). The HEK293T cell line (hereafter HEK) in which endogenous ASIC1a was removed by CRISPR/Cas9 (hereafter HEK ASIC1a^-/-^) was previously described in Borg et al. 2020^23^. HEK cells were grown in monolayer in T25, T75 or T175 flasks (Orange Scientific). The culture medium was Dulbecco’s modified Eagle’s medium (DMEM) (Thermo Fisher Scientific) supplemented with 10% fetal bovine serum (Thermo Fisher Scientific) and 1% penicillin-streptomycin (10,000 U/ml; Thermo Fisher Scientific). Cells were incubated at 37°C in a humidified 5% CO_2_ atmosphere. The cells were passaged when reaching 80% confluency and used for experiments between passage 6 and 20. HEK cell cultures were mycoplasma-tested using the standard services from Eurofins genomics.

#### Human glioblastoma stem cell line

Immortalized glioblastoma stem cell (GSC) lines, R54 and R54 ASIC1a KO were a gift from Prof. Stefan Gründer, University Hospital Aachen, Germany, and was originally isolated in the lab of Christoph Beier.^16,34^ The suspension R54 cell cultures were grown in DMEMF12, containing HEPES and NaHCO_3_ (Gibco, Thermo Fisher Scientific), supplemented with 20% B27 supplement (Gibco, Thermo Fisher Scientific), 1% glutamine (Thermo Fisher Scientific), 1% MEM vitamin (Thermo Fisher Scientific), 1% penicillin-streptomycin (Thermo Fisher Scientific), 0.1% fibroblast growth factor (FGF) and 0.1% epidermal growth factor (EGF) (Miltenyi Biotec) at 37 °C in humidified atmosphere with 5% CO_2_. The cells were seeded in concentrations of 2×10^4^ cells and passaged once a week. Glioblastoma R54 cell cultures were mycoplasma-tested using the standard services from Eurofins genomics.

### Protein purification for mass spectrometry (MS)

eGFP was used as pull-down handle on all constructs and was directly fused via a short nine residue Gly-Ala-Ser linker at the C-terminus of human ASIC1a and 10 residue Gly-Ser linker at C-terminus of human P2×7 receptor. MS sample preparation was performed in triplicate with the human P2×7-eGFP and eGFP as control constructs to determine specific interaction partners of human ASIC1a. 2×10^6^ HEK ASIC1a^-/-^ cells were seeded in 9 × 10 cm culture dishes (VWR) on day 1. On day 2, the cells were transfected with 4 ug of ASIC1a-eGFP, P2×7-eGFP or eGFP using LipoD293 (Tebu-bio) in a 1:3 ratio. On day 4, the cells were carefully washed in ice cold phosphate-buffered saline (PBS) containing Ca^2+^ and Mg^2+^ (PBS-CM), collected and lysed by adding 500 uL ice-cold lysis buffer (50 mM Tris pH 8.5, 5 mM ethylenediamine tetra acetic acid, 150 mM NaCl and 10 mM KCl, 1 μg/ml leupeptin, aprotinin and pepstatin, 1 mM phenylmethylsulfonyl fluoride, 5 mM sodium fluoride, 5 mM β-glycerophosphate and 1 mM sodium orthovanadate, 1% NP-40, 1% sodium deoxycholate, and protease and phosphatase inhibitors (complete protease inhibitor cocktail) (Roche) to each dish and collected in 1.5 mL eppendorf tubes and incubated rotating at 4°C for two hours. The lysate was centrifuged at 10,000 g at 4 °C for 10 minutes to separate the debris and soluble fraction. The lysate protein concentration was determined by bicinchoninic acid assay (Sigma-Aldrich; #BCA1) and diluted to 2 mg/mL. The lysate from each replicate was added to 25 uL of PBS-washed ChromoTek GFP-Trap® magnetic Agarose beads (Proteintech Group, Inc.), and incubated for 2 hours rotating in 4°C. Following incubation, the beads were washed four times in lysis buffer, the third wash was incubated for 30 minutes rotating at 4°C. Recovering of immunoprecipitated (IP) samples was achieved by adding 55 uL elution buffer (2X sample loading dye with 100 mM DTT) and incubated for 10 minutes at 70°C.

### MS sample preparation

The IP samples were separated using an SDS-PAGE, and each lane was cut into three pieces. Gel pieces were treated using an in-gel trypsin digestion protocol as previously described by Maurya et al. 2023 section 10^54^. In brief, Coomassie-stained SDS-PAGE bands were minced, destained, reduced,and alkylated before overnight digestion with sequencing-grade trypsin (Promega) at 37 °C. Trifluoroacetic acid (TFA) acidification was used to stop the digestion, and peptides were extracted using acetonitrile (ACN)/water. Peptides were then desalted and concentrated on C18 STAGE tips and eluted in 40% ACN, 0.5% acetic acid, and vacuum centrifuged for removing organic solvents.

### MS measurements and analysis

Peptide analysis was performed using online reversed-phase liquid chromatography coupled to a Q-Exactive HF-X quadrupole Orbitrap mass spectrometer (LC–MS/MS, Thermo Electron), as previously described^54^. Peptides were resuspended in 5% ACN, 0.1% TFA and auto-sampled into a nanoflow Easy-nLC system (Proxeon Biosystems). Separation was performed on a 15 cm self-packed PicoFrit column (75 µm internal diameter) packed with ReproSil-Pur C18-AQ 1.9 μm resin (Dr. Maisch GmbH). A 30-min multi-step gradient was utilized, from 10% to 30% Buffer B within 25 min, to 45% Buffer B within 5 min, and lastly 80% Buffer B within 30 s. Ionized peptides were examined by data-dependent acquisition using a Top12 technique. Orbitrap full-MS spectra (350–1,400 m/z) were recorded at 60,000 resolution, and precursor ions with maximum intensity were interrogated with higher-energy collisional dissociation and MS/MS spectra were recorded at 15,000 resolution. Raw MS data were analyzed by MaxQuant (version 1.6.2.3)^55^ and protein identification was performed by using the in-built Andromeda search engine against a SwissProt human protein database. Search parameters included carbamidomethyl-cysteine as static modification, and methionine oxidation, protein N-terminal acetylation, Gln→pyro-Glu conversion, and phospho(STY) as variable modifications. Two missed cleavages and six variable modifications were permitted, and peptide length was at least seven amino acids. Confidence in identification was ensured with a 1% false discovery rate (FDR) level at peptide, protein, and site decoy levels.

### MS data analysis

Label-free quantification (LFQ) data from MaxQuant output were further processed with the Perseus software package (version 1.6.2.1)^56^. Random sampling from a left-shifted normal distribution was used to impute missing values. Volcano plots were generated in Perseus, and the data were filtered to retain proteins identified in two of three replicates. Statistical significance was determined using a two-sided t-test with FDR threshold 0.05 or 0.01 and s0 > 2 trading off fold-change differences and p-value significance.

### Co-Immunoprecipitation for Western blot

2 million HEK ASIC1a^-/-^ cells were seeded into a 10 cm dish (VWR) and incubated at 37 °C in humidified 5% CO_2_ atmosphere for 24 hrs prior transfection with 4 μg ASIC and/or 12 μg MCT in combination or alone using Polyethylenimine (PEI) 25K (Polysciences) in 1:3 ratio. The same amount of cDNA was used for all ASIC and MCT subtypes, as well as for truncated MCT constructs. Six hours after transfection, cell medium was replaced with supplemented DMEM.

Cells were washed with PBS (pH 7.4) 48 hours after transfection and dislodged using cell scrapers (Orange Scientific) and transferred to a 1.5 mL eppendorf tube. After centrifugation (1,000 rpm, 5 minutes), the supernatant was aspirated, and the cell pellet was washed additionally two times with PBS. The washed cells were lysed in 0.5 mL solubilization buffer (50 mM Tris-HCl, 145 mM NaCl, 5 mM EDTA, 2 mM DDM, pH 7.5) supplemented with cOmplete EDTA-free protease inhibitor cocktail (Sigma-Aldrich) for 2 hours at 4°C and centrifuged for 30 minutes (10,000 g/4°C). In parallel, 40 μL Dynabeads Protein G (Thermo Fisher Scientific) were washed with 200 μL PBS/0.2 mM DDM and incubated with 4 μg RHO 1D4 antibody (University of British Columbia) in 50 μL PBS/0.2 mM DDM on a ferris wheel (VWR) for 30 minutes. After washing the beads with 200 μL PBS/0.2 mM DDM (with 200 μL), the cell lysate was incubated with the beads on a ferris wheel (4°C) for 90 minutes. Beads were washed with 200 μL PBS 3 times to remove nonspecifically bound proteins and incubated in 25 μL elution buffer (2:1 mixture between 50 mM glycine, pH 2.8 and 62.5 mM Tris-HCl, 2.5% SDS, 10% Glycerol, pH 6.8) supplemented with 80 mM DTT at 70°C for 10 minutes. Protein samples (12 μL) were mixed with 3 μL 5 M DTT and 5 μL 4x NuPAGE LDS sample buffer (Thermo Fisher Scientific) and incubated (95°C, 20 minutes) before SDS-PAGE using 3% to 8% TrisAcetate protein gels (Thermo Fisher Scientific). After transfer onto PVDF membranes (iBlot 2 Dry Blotting System, Thermo Fisher Scientific) and blocking in TBST with 3% nonfat dry milk for 1 hour, ASIC1a was detected using RHO 1D4 antibody (1 μg/μL, University of British Columbia) and 1:5,000 goat anti-mouse IgG HRP-conjugate (Thermo Fisher Scientific). All co-immunoprecipitation experiments were repeated two to three times each.

### [^3^H]HOCPCA uptake assays

#### Time-dependent uptake assay in *Xenopus laevis* oocytes

Oocytes were surgically removed from adult female *X. laevis* and prepared as previously described^57^. Oocytes were injected with 23 nL of RNA coding for MCT1 (1000 ng/uL) or water in equivalent volumes as a control and incubated for 3 days at 18°C before the same oocytes were injected with 23 nL of ASIC1a (100 ng/uL) or water. Injected oocytes were stored at 18°C in ND96 solution (in mM): NaCl 96, KCl 2, CaCl_2_ 1, MgCl_2_ 1, HEPES 5, pH 7.4, and supplemented with 100 IU/ml penicillin, 100 mg/ml streptomycin, pH 7.4. Transport activity was assayed 24 hrs after injections of RNA coding for ASIC1a. Uptake was measured as described previously^29^. Time-dependent uptake was measured by incubating oocytes in 2 mL Eppendorf tubes in uptake buffer (in mM: NaCl 75, KCl 2, CaCl_2_ 1, MgCl_2_ 0.82, HEPES 11, adjusted to pH 6) containing 100 nM [^3^H]HOCPCA in time-intervals between 1 and 25 minutes in room temperature (eight oocytes per condition; 100 µl total volume) Uptake was terminated by adding 1 ml of ice-cold uptake buffer to each Eppendorf tube, and oocytes were quickly transferred to Corning Net-wells inserts (Sigma-Aldrich) for washing. The oocytes were quickly washed three times with fresh ice-cold uptake buffer. Intact individual oocytes were then carefully placed in a 24-well PicoPlate (PerkinElmer), and excess buffer was carefully removed. Oocytes were then lysed by the addition of 50 µl of 2% SDS and vigorously shaken for 5–10 minutes before the addition of 450 µl of MicroScint 20 scintillation liquid (PerkinElmer), shaking for 1 hour, and quantification of radioactivity (CPM values) on a TopCount NXT reader(PerkinElmer). Data analysis was carried out using non-linear regression with one phase association exponential to give the halftimes to maximum using GraphPad Prism (10.3.1). Data are given as the means ± SD of four independent experiments.

#### Time-dependent uptake using HEK cells

The time-dependent uptake on endogenous MCT1 (in the absence and presence of endogenous ASIC1a) in HEK cells was carried out as described above. HEK cells were plated directly in poly-D-lysine (PDL)-coated (Thermo fischer scientific) white 96 well polystyrene cell culture plates (PerkinElmer), by adding 10^5^ cells per well the day before performing the uptake experiment and incubated at 37°C in humidified 5% CO_2_ atmosphere. Uptake buffer was Hanks’ Balanced Salt solution (HBSS) including 1 mM CaCl_2_, 1 mM MgCl_2_ and 20 mM HEPES, adjusted to pH 6. On the day of the experiment, the cells were carefully washed in PBS-CM before adding 100 nM [^3^H]HOCPCA uptake buffer in time-intervals between 1 and 25 minutes at room temperature (three wells per condition; 100 µl total volume). Incubation was terminated by discarding the solutions and dapping dry on paper towel, followed by washing three times with ice cold uptake buffer using a multi-channel pipette. Cells were lysed by adding 150 µL of MicroscintTM 20 scintillation liquid (PerkinElmer) to each well, shaking for 1 hour, and quantification of radioactivity (CPM values) on a TopCount NXT reader (PerkinElmer). Data analysis was carried out using non-linear regression with one phase association exponential to give the halftimes to maximum using GraphPad Prism (10.3.1). Data are given as the means ± SD of triplicate measurements of three independent experiments.

#### Saturation uptake using HEK cells

Concentration dependent uptake of [^3^H]HOCPCA was carried out in a similar fashion to the protocol described above^29^, however, using HBSS with 20 mM PIPES (pH 6) instead of HEPES. On the day of the assay, the cells were incubated with [^3^H]HOCPCA up to 100 nM and then isotope-diluted with unlabeled HOCPCA, incubated for 3 min at room temperature, and the reaction was terminated by three rapid washes with ice-cold buffer. Scintillation counting of radioactivity was performed as above, however, DPM values were obtained to allow for conversion to molar values. Total protein concentrations were determined using the Bio-Rad protein assay according to the manufacturer’s protocol to account for potential differences in the growth rate of the two cell lines. This was achieved by plating six wells per cell line in a separate clear 96-well plate at the time of plating out the cells for uptake. Data analysis was carried out using the Michaelis-Menten equation: Y = Vmax X/(Km + X), where Y is the uptake velocity, X is the substrate concentration, Vmax is the maximum uptake velocity in the same units as Y, and Km is the Michaelis−Menten constant in the same unit as X. Data were corrected for total protein and normalised to WT HEK. Data are given as the means ± SD of triplicate measurements of four independent experiments.

### KO of MCT1 in R54 cells

#### CRISPR/Cas9

CRISPR/Cas9 was used to knock out MCT1 in R54 cells. This was performed by the Transgenic Core Facility, University of Copenhagen. In short, exon 2 of the MCT1 genomic locus (first translated region) was targeted in order to generate a reading-frameshift disrupting the sequence and following expression of the gene. R54 cells were transfected with the designed guide RNA targeting exon 2 of MCT1 together with CRISR/Cas9 and individual subcloned cells were grown, genotyped by PCR and characterized by sequencing. Identified MCT1 KO clones were expanded.

#### Verification of MCT1 KO R54 by immunoprecipitation and western blot

Immunoprecipitation and western blot was performed for additional verification of MCT1 KO in R54. 2 × 10^6^ R54 cells grown in T75 flasks at 37 °C in a humidified 5% CO_2_ atmosphere was collected in 15 mL tubes and centrifuged for 2 minutes at 1,000 rpm, the supernatant was aspirated, and the cell pellet was washed additionally two times with PBS. The washed cells were lysed in 0.5 mL solubilization buffer (50 mM Tris-HCl, 145 mM NaCl, 5 mM EDTA, 2 mM DDM, pH 7.5) supplemented with cOmplete EDTA free protease inhibitor cocktail (Sigma-Aldrich) for 2 hours at 4°C and centrifuged for 30 minutes (10,000 g/4°C), and the supernatant was collected in 1.5 mL eppendorphs. The protein concentration of the lysate was determined by bicinchoninic acid assay (Sigma-Aldrich; #BCA1) and found to be between 2 to 4 mg/mL. In parallel, 40 μL Dynabeads Protein G (Thermo Fisher Scientific) were washed with 200 μL PBS, followed by incubation with 4 μg MCT1 antibody (sc-365501, Santa Cruz Biotechnology) in 50 μL PBS and rotated on a ferris wheel (VWR) for 30 minutes. After washing the beads with 200 μL PBS, the cell lysate was incubated with the beads on a ferris wheel (4°C) 90 minutes. Beads were washed with 400 μL PBS 3 times to remove nonspecific protein binding and incubated in 25 μL elution buffer (62.5 mM Tris-HCl, 2.5% SDS, 10% Glycerol, pH 6.8) supplemented with 75 mM DTT at 70°C for 10 minutes. Protein samples (14 μL) were mixed with 1 μL 1 M DTT and 5 μL 4x NuPAGE LDS sample buffer (Thermo Fisher Scientific) and incubated (95°C, 5 minutes) before SDS-PAGE using 3% to 8% TrisAcetate protein gels (Thermo Fisher Scientific). Protein bands were then transferred onto PVDF membranes (iBlot 2 Dry Blotting System, Thermo Fisher Scientific) followed by 1 hour blocking in Licor blocking buffer (TBS with 4.5 g/L Fish gelatin, 1g/L Casein, 0.02% NaN_3_). MCT1 was detected by overnight incubation with MCT1 Ab (sc-365501, Santa Cruz Biotechnology, 1:1000 dilution in blocking buffer) at 4°C, followed by 5×2 minutes washing in TBST and 1 hour incubation with anti-mouse 680 IRDye (1:5.000) (Li-Cor) in Licor blocking buffer followed by washing 5×2 minutes with TBST before imaging the blot on Syngene imager. The immunoprecipitation experiment was repeated three times.

### Patch clamp electrophysiology

#### HEK cells

HEK ASIC1a^-/-^ cells were cultured in T75 flasks to a confluency of 80-90% before splitting 1:5 into T25 flasks. Cells were incubated for 24 hours at 37°C in humidified 5% CO_2_ before the cells were transfected with 2 ug cDNA coding for ASIC1 alone or with 2 ug cDNA coding for MCT1 using Polyethylenimine (PEI) 25K (Polysciences) in 1:3 ratio. Cells were used for patch-clamp experiments 24 to 36 hrs after transfection. To ensure that the cells could be uplifted and positioned in front of a perfusion tool utilized for fast perfusion patch clamp experiments, the cells were replated onto 35-mm cell culture dishes with glass coverslips on the day of the patch clamp experiments. The transfected cells were voltage-clamped at -40 mV at room temperature in the whole-cell configuration using an Axopatch 200B amplifier (Molecular Devices) and the extracellular solution contained (in mM): NaCl 150, KCl 5, MgCl_2_ 1, CaCl_2_ 2, HEPES 10, D-glucose 10. Solutions were pH-adjusted with NaOH and HCl. Patch pipettes of borosilicate glass capillaries (World Precision Instruments) with a resistance between 3.0 to 5.0 MΩ and filled with intracellular solution containing (in mM): NaCl 30, KCl 120, MgCl_2_ 1, 0.5 CaCl_2_, EGTA 5, Na_2_ATP 2 and HEPES 10, and adjusted to pH 7.4.

#### R54 line

R54 cell cultures were passaged and seeded on PDL (Thermo fischer scientific) treated coverslips on the day of experiment and incubated at 37°C in humidified 5% CO_2_ for 1-2 hrs before conducting patch clamp experiments. The cells were voltage-clamped at -80 mV in the whole-cell configuration using an Axopatch 200B amplifier (Molecular Devices) and the extracellular solution contained (in mM): NaCl 128, KCl 5.4, HEPES 10, glucose 5.5, MgCl_2_ 1, CaCl_2_ 2; pH was adjusted to 7.4. Solutions were pH adjusted with NaOH and HCl. Patch pipettes of borosilicate glass capillaries (World Precision Instruments) with a resistance between 3.0 to 5.0 MΩ were filled with intracellular solution containing (in mM): NaCl 10, KCl 121, HEPES 10, EGTA 5, MgCl_2_ 2; pH was adjusted to 7.2. For rapid solution exchange, a custom-built glass perfusion tool equipped with four adjacent barrels, controlled by a piezo element, was used. All recordings were performed using the pCLAMP 10 software (Molecular Devices) and an Axon Digidata 1550A digitizer (Molecular Devices). Data were filtered at 10 kHz with low-pass filter, and stored continuously on a computer hard disc and analysed using pCLAMP software. Rate constants for desensitization were analysed by fitting single exponential functions to the late phase of the current trace as it returns to baseline. Statistical significance was assessed using an unpaired t-test. Data are given as the means ± SD.

### Glioblastoma RNA sequencing analysis

‘Core’ and ‘extended’ versions of the ‘GBmap’ single-cell RNA sequencing atlas from ^33^ were downloaded from the CELLxGENE service^58^ and analysed with Scanpy^59^. RNA transcript expression from the core GBmap was either displayed as the mean expression by cell type normalised to the maximum expression for a given gene, or the transcript expression for a subset of cells from each cell type was displayed separately, normalised in the same manner. The subsets were either randomly sampled when there were more than 250 cells of that type, or all cells of that type were included. To calculate the proportions of cells expressing ASIC1 and/or MCT, cells were considered as expressing if more than 0 transcripts were detected.

### Metabolism mapping

#### R54 cells

Medium was removed by collecting the R54 cells in 15 mL tubes, followed by centrifugation for 2 minutes at 1,000 rpm. The supernatant was aspirated and the cells washed and resuspended in 1 mL PBS (37°C). To enable comparison between absence and presence of ASIC1a/MCT1 same amount of cells were used within a single experiment. The amount of cells in each tube was counted in an automated cell counter (Eve™, NanoEntek) and adjusted to between 4×10^5^ to 10^6^ cells. The cells were added to 15 mL tubes and washed a second time by adding 1 mL of PBS (37°C) followed by a second spin for 2 minutes at 1,000 rpm and aspiration of the PBS. Incubation medium (2 mL) (DMEM without Glucose or Glutamine) (Sigma Aldrich) containing 10 mM L-[U-^13^C]-lactate (Cambridge Isotopes Laboratories) was added to each 15 mL tube and incubated for 30 minutes at 37°C in humidified 5% CO_2_. The uptake was terminated by spinning down the cells for 2 min at 1,000 rpm, followed by aspiration of incubation medium and addition of 2 mL ice cold PBS to each 15 mL tube, while keeping the tubes on ice to stop the metabolism of the cells. The cells were transferred to Eppendorf tubes and stored at -80°C stored until extraction. The uptake and metabolism were measured at pH 7.4 and pH 6 and each experiment was performed in two replicates per condition and repeated three times on individual days. Statistical significance was assessed using an unpaired t-test. Data are given as the means ± SD.

#### HEK cells

HEK cells were plated in 6 well plates one day prior the experiment. On the day of the experiment, the cell medium was aspirated, and the cells were washed in 1 mL PBS (37°C). The PBS was removed by aspiration before incubating the cells with 2 mL of incubation medium (DMEM without Glucose or Glutamine) (Sigma Aldrich) containing 10 mM L-[U-^13^C]-lactate (Cambridge Isotopes Laboratories) for 30 minutes. The experiment was terminated by aspiration of the incubation medium and carefully washing with 2 mL ice cold PBS. PBS was aspirated and the cells were kept in the 6 well plate and stored -80°C until extraction. The experiments were performed in pH 6 using three wells per condition (three replicates/condition) and repeated three times on individual days. Statistical significance was assessed using an unpaired t-test. Data are given as the means ± SD.

The HEK cells and R54 cells were kept on ice and extracted using 500 µl ice cold 70% ethanol and centrifuged at 4°C and 20,000 g for 20 minutes. To allow EtOH to evaporate, the cell pellet was left at room temperature with the lid open for 1 hour before adding 250 µl of 1M KOH and incubated at room temperature overnight with lids closed (to resuspend the protein). The protein suspension was stored at -20ºC until protein determination was performed by Gas-chromatography-mass-spectrometry (GC-MS).

### Mouse Experiments and Intracranial injections

Equal numbers of male and female animals were used throughout the study without any bias. Animal husbandry, ethical handling of mice and all animal work were carried out according to guidelines approved by Canadian Council on Animal Care and under protocols approved by the Toronto Centre for Phenogenomics Animal Care Committee (18-0272H). NSG mice used for xenograft experiments were NOD.Cg-Prkdcscid Il2rgtm1Wjl/SzJ mice [Jackson laboratories #005557]. R54 WT, R54 MCT KO, and R54 ASIC1a KO cells were diluted to a final concentration of 100,000 cells/uL, mixed with 0.05% Fast Green (F7252-5G) and loaded into a syringe (Hamilton 7659-01) with 33-gauge needle (Hamilton 7803-05). P0 pups were anesthetized on parafilm covered ice and their head secured with a custom 3D-printed mold. A stereotactic manipulator was used to position the needle to 0.3 mm above the Bregma towards the Lambda Suture and 0.1 mm lateral of Sagittal Suture into the right ventricle. The needle punctured 3 mm into the skull, and retracted 1mm for a final depth of 2 mm. 1 ul of cells (50,000 cells) was administered and allowed 1 minute to diffuse before retraction of the needle. Post-injection, the neonates were warmed on a heating pad. A total of 15 mice were included in the experiment and statistical significance was assessed using Mantel-Cox Test.

