## Supplementary Information for "Coupling between acid-sensing ion channel 1a and the monocarboxylate transporter family shapes cellular pH response"

Figure S1:ASIC1a-MCT1 coupling modulates ASIC1a function — Page 2

Figure S2: Functional modulation of ASIC1a is independent of MCT1 conformation — Page 3

Figure S3: ASIC1a has no major impact on MCT1 transport activity — Page 4

Figure S4: Peripheral MCT helices are essential for regulating ASIC1a function — Page 5

Figure S5: MCT1 selectively modulates ASIC1a — Page 6

Figure S6: High expression of MCT1 and ASIC1a coding RNA in human glioblastoma — Page 7

Table S1: Desensitization kinetics and pH sensitivity — Page 8

Table S2: Construct overview — Page 9

Table S3: Chimeric and truncated constructs — Page 10

References: — Page 11

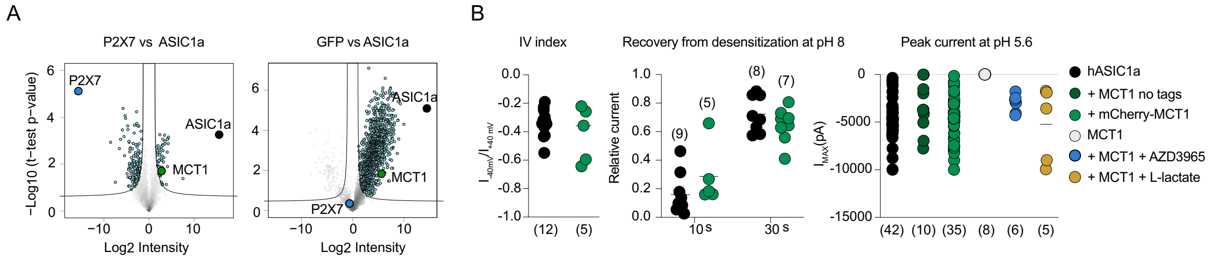


**Figure S1. ASIC1a-MCT1 coupling modulates ASIC1a function. A** Volcano plots showing proteins (light blue circles) identified by MS following co-immunoprecipitation using ASIC1a-GFP (black circles), P2X7-GFP (blue circles) or GFP as bait. Left plot; illustrating hP2X7-GFP and potential interaction partners (left) (as a control) and ASIC1-GFP and potential interaction partners (right). Right graph; illustrating GFP and potential interaction partners (as a control) and ASIC1a and potential interaction partners (right). **B** MCT1 has no effect on ASIC1a IV index (relative current at -40mV and 40mV) (left) and recovery after 10 and 30 s in pH 8 (middle). ASIC1a mediated current amplitude at pH 5.6 (right) was not affected by mCherry-MCT1 (mCherry tag for visualization) (green circles) or without mCherry tag (dark green circles). MCT1 alone (grey circles) did not give rise to currents by pH 5.6. Addition of AZD3965 (blue circles) or L-lactate (yellow circles) during patch clamp recordings on HEK ASIC1a $^{-/-}$ cells overexpressing ASIC1a and MCT1 had no effect on current amplitude at pH 5.6 (right). Number of repetitions in brackets.

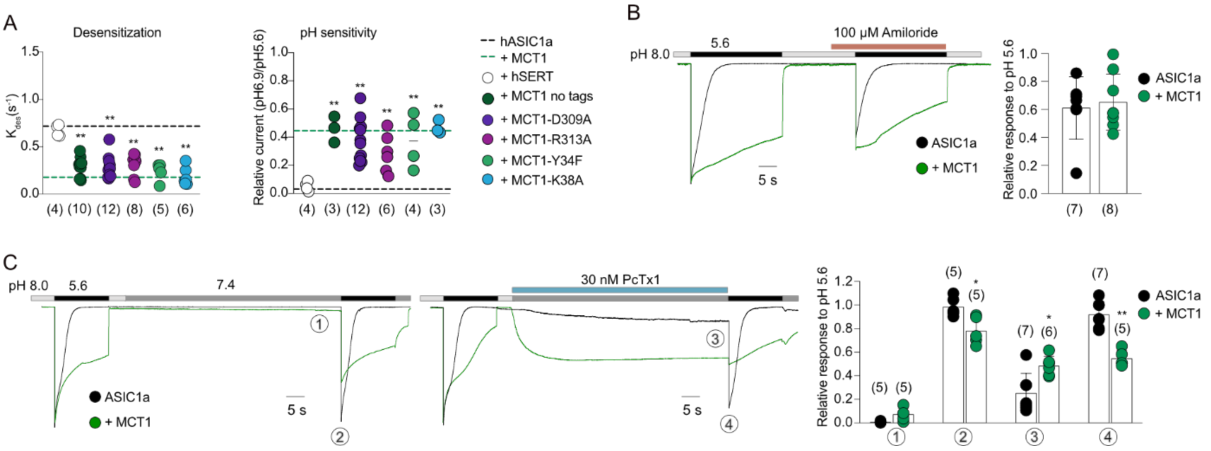


**Figure S2. Functional modulation of ASIC1a is independent of MCT1 conformation, while ASIC1a-MCT1 coupling affects ASIC1a pharmacology.** **A** Summary of desensitization kinetics (left) and pH sensitivity (right) from HEK ASIC1a$^{-/-}$ cells overexpressing ASIC1a alone (average illustrated by black dashed line), with mCherry-MCT1 (average illustrated by green dashed line), with human serotonin transporter (SERT) as a control, MCT1 without mCherry (dark green circles), and with MCT1-mutants neutralizing the L-lactate and proton binding sites. Asterisks denote significance levels in comparison to ASIC1a alone: $^{**}p < 0.001$, unpaired t-test. Number of repetitions in brackets. **B** Current trace illustrating whole cell patch clamp recordings from HEK ASIC1a$^{-/-}$ cells overexpressing ASIC1a alone (black trace) and with MCT1 (green trace) (left) in absence and presence of 100 µM amiloride. Graph; summary of relative currents at pH 5.6 before and after application of 100 µM amiloride. Number of repetitions in brackets. **C** Current trace illustrating whole cell patch clamp recordings from HEK ASIC1a$^{-/-}$ cells overexpressing ASIC1a alone (black trace) and with MCT1 (green trace) (left) and in presence of 30 nM PcTx1 (right). Graph; summary of relative ASIC1a currents under various conditions: (1) at pH 7.4 without (black circles) and with MCT1 (green circles), (2) at pH 5.6 following wash at pH 7.4 without (black circles) and with MCT1 (green circles), (3) at pH 7.4 with 30 nM PcTx1 without (black circles) and with MCT1 (green circles), and (4) at pH 5.6 following wash at pH 7.4 with 30 nM PcTx1 without (black circles) and with MCT1 (green circles). Asterisks indicate significance levels relative to ASIC1a alone: $^{*}p < 0.05$ and $^{**}p < 0.001$, unpaired t-test. Number of repetitions in brackets.

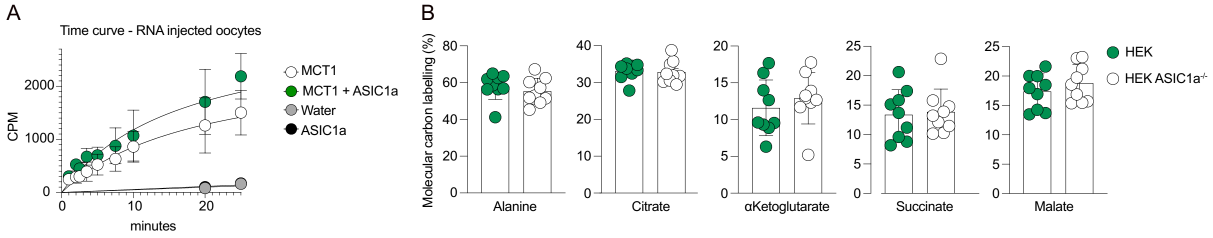


**Figure S3. ASIC1a has no major impact on MCT1 transport activity. A** Graphs illustrating uptake of [$^3$H]HOCPCA radioligand (substitute for lactate) in counts pr minute (CPM) measured over time using oocytes injected with RNA coding for MCT1 alone (white circles), MCT1 and ASIC1a (green circles), ASIC1a alone (black circles) or water (grey circles). MCT1 alone or in presence of ASIC1a showed half-time to maximum at similar values (13.63 ± 13.19 min (22) and 6.20 ± 5.19 min (19), respectively ($p>0.05$, unpaired t-test)). The experiment was repeated on four different batches of oocytes on four individual days, with the number in brackets illustrating total number of oocytes included and error bars as standard deviation. **B** Graphs showing total $^{13}$C-carbon incorporation in TCA cycle metabolites alanine, citrate alpha-ketoglutarate, succinate and malate following uptake of $^{13}$C-labbelled L-lactate via endogenous MCT1 in presence (green circles) and absence of endogenous ASIC1a (white circles) using HEK WT and HEK ASIC1a$^{-/-}$ cells. Three replicates/condition was repeated on three individual experiment days.

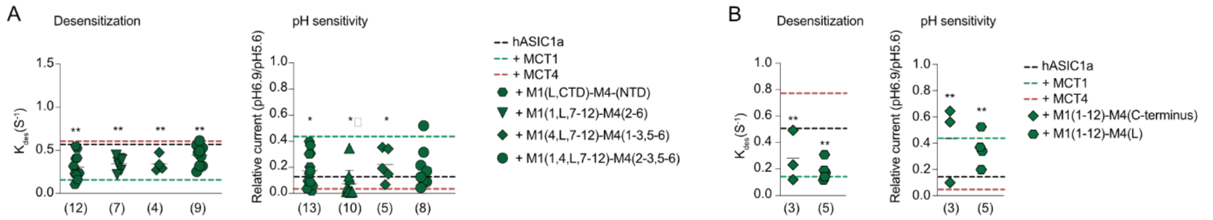


**Figure S4. Peripheral MCT helices are essential for regulating ASIC1a function.** **A** Summary of desensitization kinetics at pH 5.6 (left) and pH sensitivity (right) of ASIC1a alone (black dotted line), with MCT1 (green dotted line) and MCT4 (red dotted line) as well as ASIC1a with MCT1-MCT4 chimeric constructs retaining additional MCT1 helices, along with the CTD of MCT1 (green symbols) **B** The intracellular C-terminus and linker are non-essential for MCT1 regulation of ASIC1a function. Summary of desensitization kinetics at pH 5.6 (left) and pH sensitivity (right) when exchanging the intracellular C-tail MCT1 with the corresponding in MCT4 (green diamonds) or the linker between MCT1 NTD and CTD with the corresponding linker in MCT4 (green hexagons). Asterisks indicate significance levels relative to ASIC1a alone: *p < 0.05 and **p < 0.001, unpaired t-test. Number of repetitions in brackets.

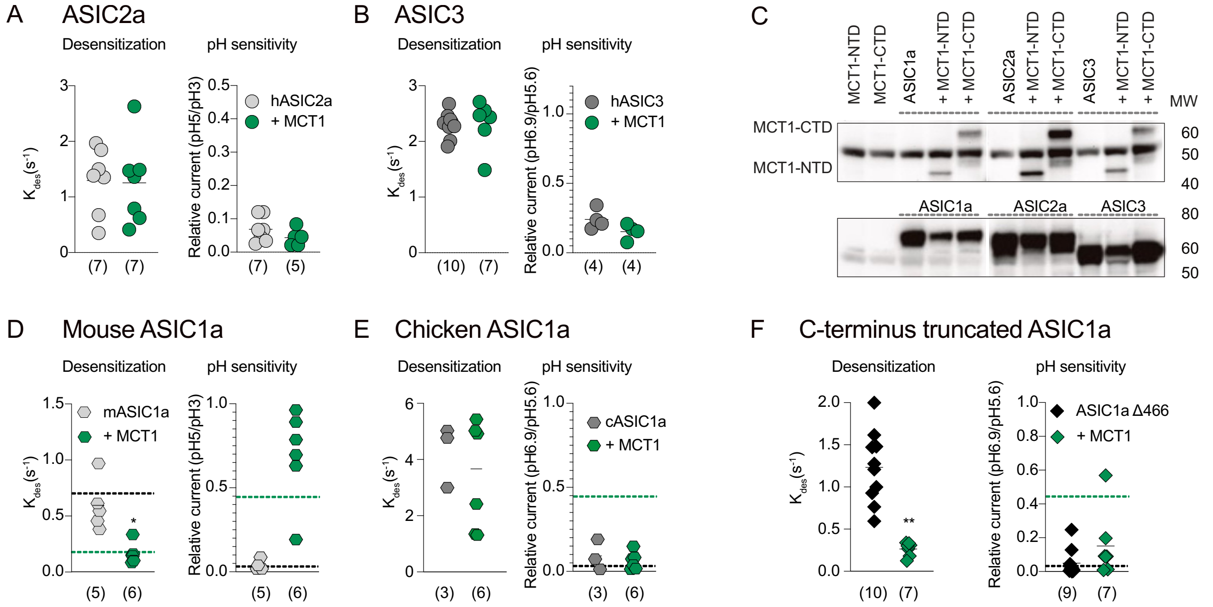


**Figure S5. MCT1 selectively modulates ASIC1a. A-B** Only ASIC1a is modulated by MCT1. Dot-plot summarizing desensitization kinetics at pH 5.6 (left) and pH sensitivity index (right) for ASIC2a (grey circles) alone and with MCT1 (green circles) (**A**) and for ASIC3 alone (dark grey circles) and with MCT1 (green circles) (**B**). Number in brackets illustrates number of repetitions. **C** Western blot illustrating the pull down of individual truncated MCT1 versions comprised of only NTD or CTD using hASIC1a, hASIC2a or hASIC3 as bait. All three ASIC subtypes can pull down the MCT1-NTD and CTD. The co-immunoprecipitation experiment was repeated two times. **D-E** Human and mouse ASIC1a share 97.9% sequence identity and are both functionally modulated by human MCT1. In contrast, chicken ASIC1a shares only 63.8% sequence identity with human ASIC1a and is not modulated by human MCT1. Summary of desensitization kinetics (left) and pH sensitivity (right) of mouse ASIC1 alone (ight grey hexagon) and with MCT1 (green hexagon) (**D**). Summary of desensitization kinetics (left) and pH sensitivity (right) of chicken ASIC1 alone (grey hexagon) and with MCT1 (green hexagon) (**E**). Asterisks denote significance levels compared to ASIC1a with MCT1 alone: * $p < 0.05$, unpaired t-test. Number in brackets illustrates number of repetitions. **F** The C-terminus of ASIC1 is not involved in coupling with MCT1. Summary of desensitization kinetics (left) and pH sensitivity (right) for ASIC1aΔ466 alone (black diamonds) and with MCT1 (green diamonds). Asterisks denote significance levels compared to ASIC1a with MCT1 alone:**$p < 0.001$, unpaired t-test. Number of repetitions in brackets.

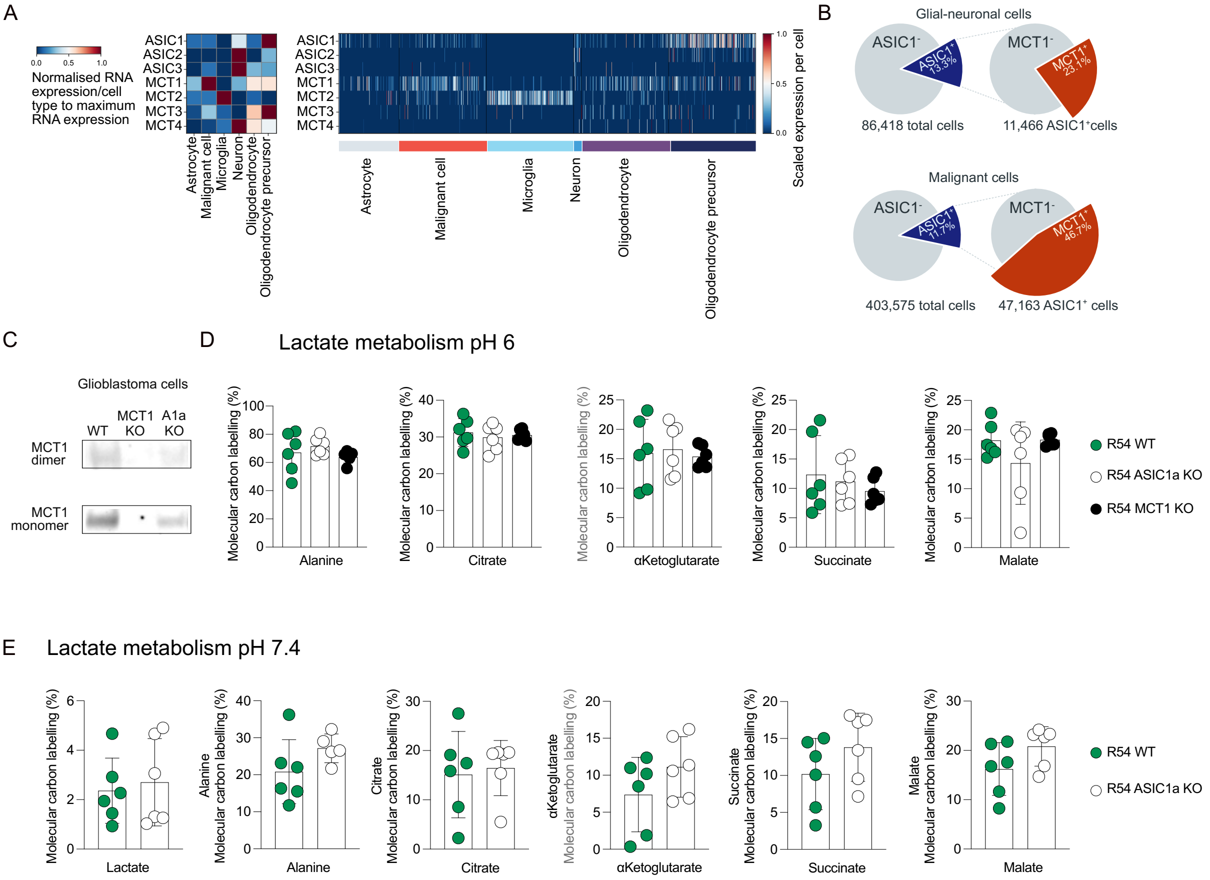


**Figure S6. High expression of MCT1 and ASIC1a coding RNA in human glioblastoma. A** The dataset of single-cell RNA sequencing from 240 patients, assembled by Ruiz-Moreno et al.[1], was used to summarize the RNA expression of ASIC and MCT subtypes in various human brain cells; astrocytes, glioblastoma (malignant cells), microglia, neurons, oligodendrocytes, and oligodendrocyte precursor cells. Left; Mean RNA expression level per cell type normalized to the maximum level of RNA expression and color-coded accordingly. Right; Scaled RNA expression per cell, subsampled to show a maximum of 250 cells per cell type, with each line representing subtype specific RNA expression. **B** Diagram based on analysis from the dataset of Ruiz-Moreno et al.[1], showing the upregulation of MCT1 RNA expression in malignant cells. **C** Western blow illustrating the CRISPR-Cas KO of MCT1 in R54 glioblastoma. **D** Graphs showing total amount of incorporation $^{13}$C -carbon in TCA cycle metabolites following $^{13}$C labelled L-lactate uptake in presence (green circles) or absence of endogenous hASIC1a (white circles) and in absence of MCT1 (black circles) in human Glioblastoma at pH 6. **E** Graphs showing total amount of $^{13}$C labelled L-lactate uptake and the following incorporation $^{13}$C -carbon in TCA cycle metabolites in presence (green circles) or absence of endogenous hASIC1a (white circles) and in absence of MCT1 (black circles) in human Glioblastoma at pH 7.4.

**Table S1. Desensitization kinetics and pH sensitivity**. Data are shown as mean ± SD, with number of repetitions in brackets. Note that all MCT constructs, with the exception of 'MCT1 no tags', were tagged with mCherry at the N-terminus for visualization during patch-clamp recordings. SERT was tagged at the C-terminus. The MCT-NTD construct consists of membrane helix 1-6 and MCT-CTD consists of membrane helix 7-12. L is the linker between helix 6 and 7 connecting the MCT-NTD and MCT-CTD.

| ASIC subtypes | Desensitization kinetics ($s^{-1}$) | pH sensitivity ($I_{pH\ 6.9}/I_{pH5.6}$) in % |
|---|---|---|
| Human ASIC1a | 0.72 ± 0.15 (36) | 3 ± 5 (35) |
| + MCT1 no tags | 0.29 ± 0.10 (10) | 46 ± 9 (3) |
| + MCT1 | 0.18 ± 0.07 (29) | 45 ±17(31) |
| + MCT2 | 0.47 ± 0.09 (5) | 5 ± 3 (5) |
| + MCT4 | 0.81 ± 0.22 (10) | 1 ± 3 (10) |
| + SERT | 0.67 ± 0.07 (4) | 5 ± 3 (4) |
| Human ASIC1a Δ466 | 1.23 ± 0.43 (10) | 6 ± 8 (9) |
| + MCT1 | 0.27 ± 0.08 (7) | 15 ± 29 (7) |
| Human ASIC1b | 5.81 ± 3.44 (13) | 0.2 ± 0.2 (13) |
| + MCT1 | 6.46 ± 2.66 (12) | 0.5 ± 0.3 (12) |
| Human ASIC2a | 1.30 ± 0.58 (7) | 7 ± 4 (7) |
| + MCT1 | 1.25 ± 0.74 (7) | 4 ± 3 (5) |
| Human ASIC3 | 2.30 ± 0.23 (10) | 24 ± 7 (4) |
| + MCT1 | 2.42 ± 0.49 (7) | 15 ± 6 (4) |
| Chicken ASIC1 | 4.26 ± 1.10 (3) | 9 ± 9 (3) |
| + MCT1 | 3.41 ± 1.93 (6) | 6 ± 5 (6) |
| Mouse ASIC1a | 0.60 ± 0.23 (5) | 4 ± 3 (5) |
| + MCT1 | 0.16 ± 0.09 (6) | 69 ± 27 (6) |
| ASIC1a & ASIC1b chimeras | | |
| ASIC1a-1b-ECD | 4.44 ± 1.28 (5) | 1 ± 1 (5) |
| + MCT1 | 1.71 ± 1.94 (8) | 9 ± 11 (8) |
| ASIC1a-1b-M1 | 0.56 ± 0.13 (5) | 4 ± 4 (5) |
| + MCT1 | 0.20 ± 0.03 (4) | 31 ± 29 (4) |
| ASIC1a-1b-NT-PreM1 | 0.45 ± 0.07 (8) | 1 ± 0.4 (8) |
| + MCT1 | 0.48 ± 0.11 (8) | 1 ± 1 (8) |
| ASIC1a + MCT1 NTD or CTD | | |
| + M1-NTD (1:1) | 0.66 ± 0.16 (5) | 2 ± 2 (7) |
| + M1-CTD (1:1) | 0.75 ± 0.15 (5) | 8 ± 8 (4) |
| + M1-CTD (1:3) | 0.67 ± 0.11 (3) | 0.1 ± 0.2 (3) |
| + M1-NTD + CTD (1:1) | 0.53 ± 0.06 (4) | 11 ± 7 (4) |
| + M1-NTD + CTD (1:3) | 0.21 ± 0.12 (4) | 40 ± 15 (4) |
| ASIC1a with MCT1 and MCT4 chimeras | | |
| + M1(NTD)-M4(L, CTD) | 1.00 ± 0.13 (4) | 0.3 ± 0.4 (5) |
| + M1(CTD)-M4(NTD, L) | 0.41 ± 0.10 (11) | 8 ± 0.12 (11) |
| + M1(L, CTD)- M4(NTD) | 0.30 ± 0.14 (12) | 17 ±13 (13) |
| + M1(1,L,7-12)-M4(2-6) | 0.35 ± 0.08 (7) | 20 ±16 (8) |
| + M1(4,L,7-12)-M4(1-3, 5-6) | 0.35 ± 0.09 (4) | 22 ±12 (5) |
| + M1(1,4, L, 7-12)-M4(2-3, 5-6) | 0.44 ± 0.12 (9) | 7 ± 11 (19) |
| + M1(1-12)-M4(L) | 0.19 ± 0.07 (5) | 36 ± 13 (4) |
| + M1(1-12)-M4(C-terminus) | 0.28 ± 0.19 (3) | 43 ±29 (3) |
| ASIC1a with MCT1 mutants | | |
| + MCT1-Y34F | 0.24 ±0.9 (5) | 37 ± 19 (4) |
| + MCT1-K38A | 0.19 ± 0.10 (6) | 47 ± 5 (3) |
| + MCT1-D309N | 0.13 ±0.07 (7) | 49 ± 20 (9) |
| + MCT1-D309A | 0.29 ± 0.11 (12) | 38 ± 15 (6) |
| + MCT1-R313A | 0.31 ±0.11 (8) | 29 ±14 (9) |
| ASIC1a with MCT1 and compounds | | |
| + AZD3965 | 0.31 ± 0.21 (6) | 45 ± 22 (6) |
| + L-lactate | 0.20 ± 0.05 (5) | 25 ± 9 (5) |

**Table S2. Construct overview**. All constructs were purchased from TWIST Bioscience and designed to contain a FLAG, 1D4 or HA tag as pull-down handle and either directly fused with mCherry or eGFP or separated by P2A/IRES. Note that MCT constructs were tagged at the N-terminus and ASIC constructs at the C-terminus.

| ASIC subtypes | Protein entry # |
|---|---|
| Human ASIC1a | P78348-2 |
| Human ASIC1b | P78348-3 |
| Human ASIC2a | Q16515-1 |
| Human ASIC3 | Q9UHC3-1 |
| Chicken ASIC1 | Q1XA76 |
| Mouse ASIC1a | Q6NXK8-1 |
| **MCT subtypes** | **Protein entry #** |
| Human MCT1 | P53985-1 |
| Human MCT2 | O60669 |
| Human MCT4 | O15427 |
| **Other constructs** | **Protein entry #** |
| Human SERT | P31645-1 |
| Human P2X7 | Q99572-1 |

**Table S3. Chimeric and truncated constructs.** All constructs were purchased from TWIST Bioscience and designed to contain a FLAG, 1D4 or HA tag as pull-down handle, either directly fused with mCherry or eGFP or separated by P2A/IRES. Note that MCT constructs were tagged at the N-terminus and ASIC constructs at the C-terminus. All chimeric constructs were generated using the original protein templates (see protein entry #) and tagged with mCherry at the N-terminus. SERT was tagged at the C-terminus. MCT1 and MCT4 is abbreviated M1 and M4, respectively. The MCT-NTD consists of membrane helix 1-6 and MCT-CTD consists of membrane helix 7-12. L, linker between helix 6 and 7 connecting the MCT-NTD and MCT-CTD.

| ASIC1a and 1b Chimeric constructs | ASIC1a | ASIC1b |
|---|---|---|
| ASIC1a-1b-NT-PreM1 | 40-528 | 1-85 |
| ASIC1a-1b-M1 | 1-40, 70-528 | 86-115 |
| ASIC1a-1b-ECD | 1-69, 186-528 | 116-219 |
| ASIC1a Truncated constructs | ASIC1a | Linker |
| ASIC1a Δ466 | 1-466 | |
| NT-PreM1-M1-M2* | 1-72, 425-466 | 16 AA of GSA |
| M1-M2* | 38-72, 425-466 | 16 AA of GSA |
| NT-PreM1 | 1-39 | - |
| NT-PreM1-M1 | 1-72 | - |
| MCT1-MCT4 Chimeric constructs | MCT1 | MCT4 |
| M1(NTD)-M4(L, CTD) | 1-259 | 225-465 |
| M1(CTD)-M4(NTD, L) | 260-500 | 1-224 |
| M1(L, CTD)- M4(NTD) | 199-500 | 1-200 |
| M1(1,7-12)- M4(2-6) | 1-53, 199-500 | 56-200 |
| M1(4,L,7-12)-M4(1-3, 5-6) | 107-139,199-500 | 1-108, 142-200 |
| M1(1,4, L, 7-12)-M4(2-3, 5-6) | 1-53,107-139,199-500 | 56-108, 142-200 |
| M1(1-12)-M4(L) | 1-187, 261-500 | 201-225 |
| M1(1-12)-M4(C-terminus) | 1-449 | 412-465 |
| MCT Truncated constructs | MCT1 | MCT4 |
| MCT1-NTD (1-6) | 1-227 | - |
| MCT1-CTD (7-12) | 228-500 | - |
| MCT4-NTD (1-6) | - | 1-201 |
| MCT4-CTD (7-12) | - | 202-465 |

*constructs has a 16 AA linker of Gly, Ser and Ala (GASGGSASAGSAGSAS ) **L is the intracellular loop between M6 and M7, this loop differs between MCT1 and MCT4.
